# Cohesin disrupts polycomb-dependent chromosome interactions

**DOI:** 10.1101/593970

**Authors:** JDP Rhodes, A Feldmann, B Hernández-Rodríguez, N Díaz, JM Brown, NA Fursova, NP Blackledge, P Prathapan, P Dobrinic, M Huseyin, A Szczurek, K Kruse, KA Nasmyth, VJ Buckle, JM Vaquerizas, RJ Klose

## Abstract

How chromosome organisation is related to genome function remains poorly understood. Cohesin, loop-extrusion, and CTCF have been proposed to create structures called topologically associating domains (TADs) to regulate gene expression. Here, we examine chromosome conformation in embryonic stem cells lacking cohesin and find as in other cell types that cohesin is required to create TADs and regulate A/B compartmentalisation. However, in the absence of cohesin we identify a series of long-range chromosomal interactions that persist. These correspond to regions of the genome occupied by the polycomb repressive system, depend on PRC1, and we discover that cohesin counteracts these interactions. This disruptive activity is independent of CTCF and TADs, and regulates gene repression by the polycomb system. Therefore, in contrast to the proposal that cohesin creates structure in chromosomes, we discover a new role for cohesin in disrupting polycomb-dependent chromosome interactions to regulate gene expression.

## Introduction

Spatial organisation of the genome influences gene transcription and other fundamental DNA-based processes. Recently, genome-wide chromosome conformation capture (Hi-C) has significantly advanced our understanding of chromosomal organisation (Rowley and Corces, 2018). This has shown that megabase-sized regions of chromosomes, which have similar transcriptional activity and chromatin modifications, tend to interact preferentially. When these interactions involve active regions of chromosomes they are referred to as A compartments and interactions between less active regions are referred to as B compartments (Lieberman-Aiden et al., 2009). At the sub-megabase scale, chromosomes are partitioned into topologically associating domains (TADs) which correspond to contiguous regions of chromatin that interact more frequently than with chromatin outside the domain (Dixon et al., 2012; Nora et al., 2012; Rao et al., 2014). There is increasing evidence that TAD formation occurs through a process called loop extrusion. It has been proposed that cohesin can utilize its ATPase activity to extrude loops of chromatin and that this is limited or terminated by CTCF occupied insulator DNA elements (Fudenberg et al., 2016; Sanborn et al., 2015). This process is thought to structure and insulate chromosomes, limiting the effects of distal gene regulatory elements to genes within a given TAD. Indeed, alterations in TAD boundaries can lead to perturbed gene expression and human disease (Lupiáñez et al., 2015). Importantly, the function of cohesin in loop extrusion appears to be distinct from its essential and well characterised role in sister chromatid cohesion (Guacci et al., 1997; Michaelis et al., 1997).

Based on these observations, super-resolution chromosome imaging has been applied to test whether the organisational concepts which emerge from ensemble Hi-C experiments are also evident in single cells (Bintu et al., 2018; Finn et al., 2019; Miron et al., 2019). This has revealed contiguous globular chromosomal structures that are independent of cohesin and loop extrusion, and spatially heterogeneous amongst individual cells. Moreover, single cell Hi-C experiments indicate that interactions within TADs are infrequent (Flyamer et al., 2017). This suggests that TADs are not static structural entities, but result from tendencies to interact, which only become evident when averaged over a population of cells in ensemble Hi-C analysis.

If TADs are not fixed structural entities, then fundamental questions remain as to what roles cohesin and loop extrusion have in regulating interphase chromosome structure and function. Recent attempts to address these questions have proposed that cohesin regulates interactions between super-enhancers in cancer cells (Rao et al., 2017) and helps to actively guide distant enhancers to their target genes in somatic cells (Hadjur et al., 2009). However, to what extent these processes function in different cell types, how they are related to CTCF/TADs, and what role they play in gene regulation remains poorly defined.

To address these questions, we removed cohesin in mouse embryonic stem cells (ESCs) and examined chromosome interactions by Hi-C. We show that cohesin loss eliminates TADs and enhances A/B compartmentalisation as in other cell types. However, in the absence of cohesin we find that a series of long-range high frequency interactions corresponding to regions of the genome occupied by the polycomb repressive complexes (PRC1 and PRC2) persist. These interactions rely on PRC1 and interestingly, in the absence of cohesin, we discover that interactions between polycomb chromatin domains are strengthened. Using single cell analysis we demonstrate that cohesin separates polycomb chromatin domains, explaining the effects observed by Hi-C. Removal of CTCF, and disruption of TADs, does not strengthen these interactions, revealing that cohesin counteracts the association of polycomb chromatin domains through mechanisms that are independent of TADs or insulation. Moreover, we find that increases in polycomb chromatin domain interactions following cohesin loss has functional consequences on gene expression. Together these discoveries reveal a new role for cohesin in disrupting polycomb dependent chromosome interactions and gene repression.

## Results

### Cohesin-independent chromosomal interactions exist in ESCs

We chose to study the loss of cohesin in ESCs because they are non-transformed, diploid, and have a wealth of existing genomic information characterising their chromosome structure and chromatin modifications. To do this we used CRISPR/Cas9-based genome engineering and developed an ESC line in which the cohesin subunit SCC1 (RAD21), could be rapidly removed via an auxin inducible degron (Natsume et al., 2016) (Figure 1A, B and S1A). This circumvented complications associated with the absence of cohesin during cell division and allowed us to examine the effects that loss of cohesin has on chromosome structure and function.

**Figure 1.**
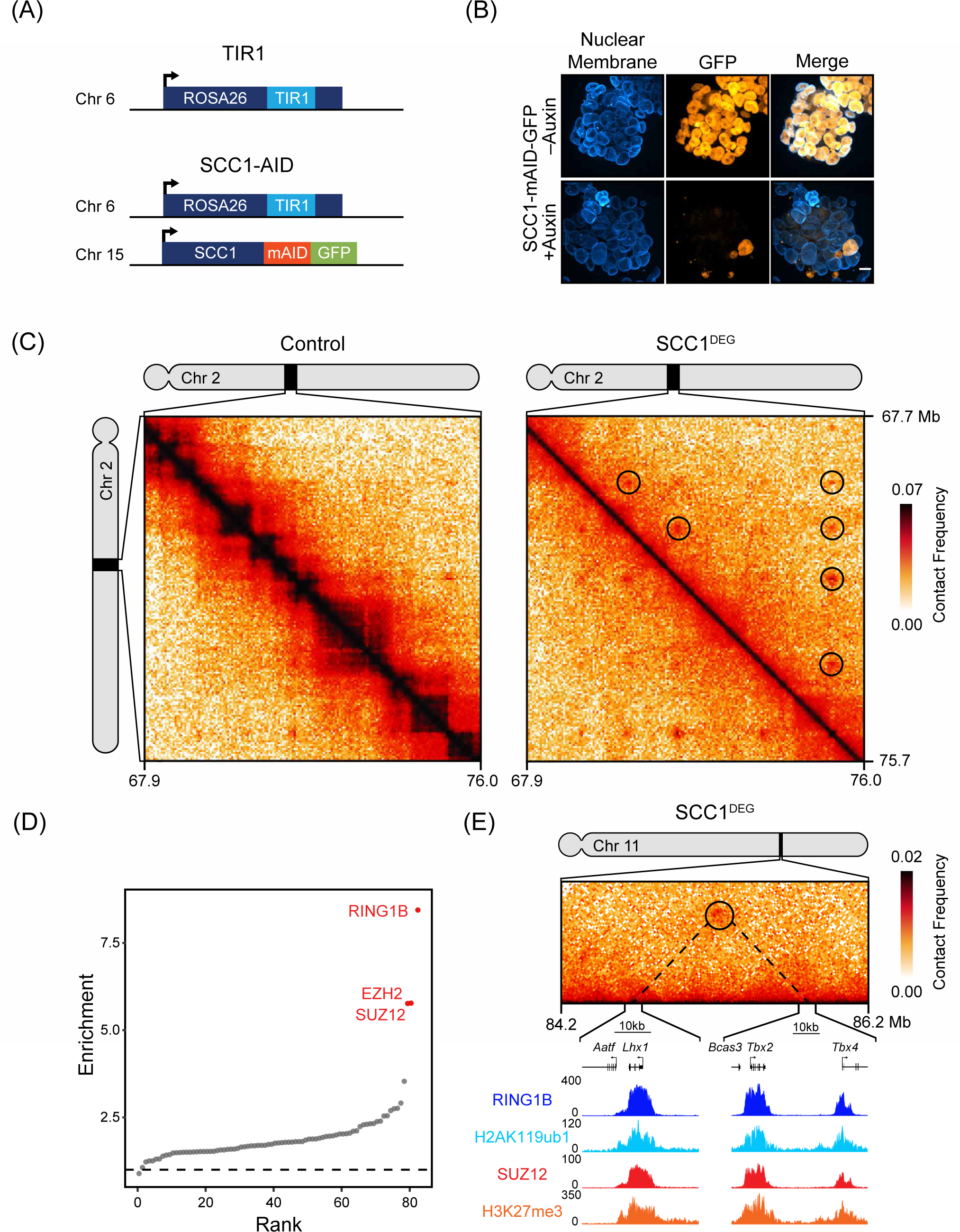
Cohesin-independent chromosomal interactions correspond to polycomb chromatin domains in ESCs. (A) A schematic illustrating the genotype of the TIR1 and SCC1-mAID-GFP cell lines developed for Hi-C. (B) Immunofluorescence microscopy images of SCC1-mAID-GFP ESCs ± auxin (6 h). The nuclear membrane was labelled with an antibody against Lamin B1. Scale bar = 10 µm (bottom). (C) Hi-C in Control (TIR1 line + auxin) (left) and SCC1^DEG^ (SCC1-mAID-GFP line + auxin) (right) cells after auxin treatment visualised at 40 kb resolution. Peaks identified on the SCC1^DEG^ Hi-C matrix are shown as black circles. The genomic co-ordinates are illustrated below and to the right of the matrices. (D) Enrichment of histone modifications and proteins at paired interaction sites compared to the enrichments at matched random interaction sites. (E) ChIP-Seq snapshot illustrating RING1B, H2AK119ub, SUZ12 and H3K27me3 under an interaction that persists in the absence of cohesin. The Hi-C matrix is shown above at 20 kb resolution.

To examine chromosome interactions in the absence of cohesin we treated cells with auxin for 6 hours to allow the effects of cohesin loss to manifest and compared *in situ* Hi-C (Díaz et al., 2018; Rao et al., 2014) matrices from the SCC1 degron ESCs (SCC1^DEG^) and control ESCs. Consistent with previous findings (Rao et al., 2017; Schwarzer et al., 2017; Wutz et al., 2017), removal of cohesin caused a complete loss of TADs (Figure S1B and C) and modestly enhanced A/B compartmentalisation (Figure S1D). However, visual inspection of the Hi-C matrices also revealed numerous interactions that were evident in control cells and persisted in the absence of cohesin (Figure 1C). We then used computational approaches to identify these persistent interactions throughout the genome (Rao et al., 2014) and uncovered 336 sites of high interaction frequency in cohesin-depleted cells. Interestingly, when we examined whether there were any DNA binding factors or chromatin features associated with these interaction sites, there was a strong enrichment of proteins that form polycomb repressive complexes (PRC1 and PRC2) (Figure 1D). This association was further evident when the occupancy of PRC1, PRC2, and their histone modifications were examined at interaction sites (Figure 1E and S1E). The most enriched polycomb protein at these sites was the PRC1 component RING1B. When we examined its occupancy in more detail, we found that 85% (287/336) of interactions had RING1B associated with at least one of the interaction sites and 65% (218/336) had RING1B at both interaction sites. Interestingly, these interactions tended to involve large polycomb chromatin domains, suggesting that the size of the domain may contribute to interaction frequency (Figure S1F). Therefore, removal of cohesin in ESCs leads to strengthening of A/B compartmentalisation and loss of TADs, but some strong chromosomal interactions persist and these correspond to regions of the chromosome occupied by the polycomb system.

### Polycomb mediates interactions that persist in the absence of cohesin

In ESCs it is known that polycomb chromatin domains can associate with each other, even over very long distances (Bonev et al., 2017; Denholtz et al., 2013; Joshi et al., 2015; Schoenfelder et al., 2015). To determine whether sites that persisted in the absence of cohesin rely on the polycomb system for their formation, an AID tag was added to RING1B and the closely related and interchangeable paralogue, RING1A, was deleted (Figure 2A, B and S2A). We then treated cells for 6 hours with auxin to remove RING1B (RING1B^DEG^) and carried out *in situ* Hi-C. Examination of genomic distance-dependent contact probabilities and TADs following removal of PRC1 revealed that these features were unaffected (Figure 2C and D) with minor increases in A/B compartmentalisation (Figure S2B). However, the interactions at sites that persisted in the absence of cohesin were lost from the Hi-C matrices (Figure 2E, F and S2C). Therefore, in ESCs, PRC1 contributes little to A/B compartmentalisation and TADs, but is responsible for long-range chromosomal interactions that also persist in the absence cohesin.

**Figure 2.**
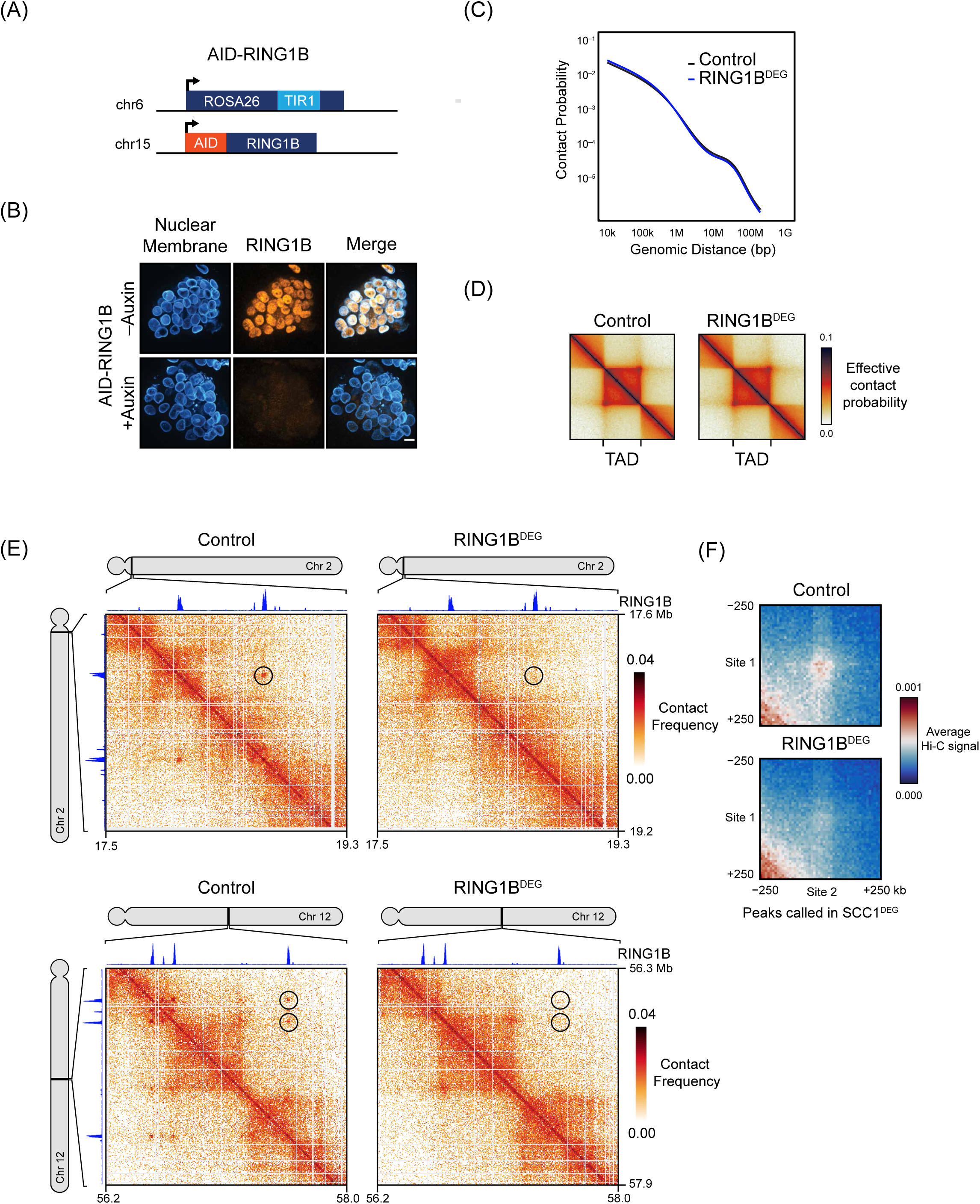
Polycomb mediates interactions that persist in the absence of cohesin. (A) A schematic illustrating the genotype of the AID-RING1B cell line. (B) Immunofluorescence microscopy images of AID-RING1B ESCs ± auxin (6 h incubation) (bottom). The cells were labelled with antibodies against Lamin B1 and RING1B. Scale bar = 10 µm (bottom). (C) Genomic distance-dependent contact probability from Hi-C in Control or RING1B^DEG^ (AID-RING1B + auxin) cells. (D) Aggregate TAD analysis of Control and RING1B^DEG^ cells at 10 kb resolution. Effective contact probability is displayed at a published set of TAD intervals from ESC Hi-C (Bonev et al., 2017). (E) Hi-C in Control and RING1B^DEG^ cells at interactions that persist in the absence of cohesin (black circles) at 5 kb resolution. RING1B ChIP-seq is displayed above and to the left of the matrices. (F) Aggregate analysis of Hi-C from Control and RING1B^DEG^ cells at interactions that persist in the absence of cohesin (n=336).

### Cohesin removal strengthens long-range polycomb chromatin domain interactions

In the absence of cohesin, we noticed that the interaction frequency between polycomb chromatin domains often appeared to increase in the Hi-C matrices, suggesting that cohesin may regulate these interactions (Figure 1C and 3A). Indeed, aggregate analysis of the interactions that persist in the absence of cohesin, and which we have shown rely on PRC1 to form, displayed a strong increase in interaction frequency (Figure 3B). Importantly, these effects did not result from increases in PRC1 occupancy, as RING1B binding was similar, or even slightly lower, than in cells with normal cohesin levels (Figure 3C and D). Together this reveals that cohesin regulates polycomb chromatin domain interactions without affecting PRC1 occupancy.

**Figure 3.**
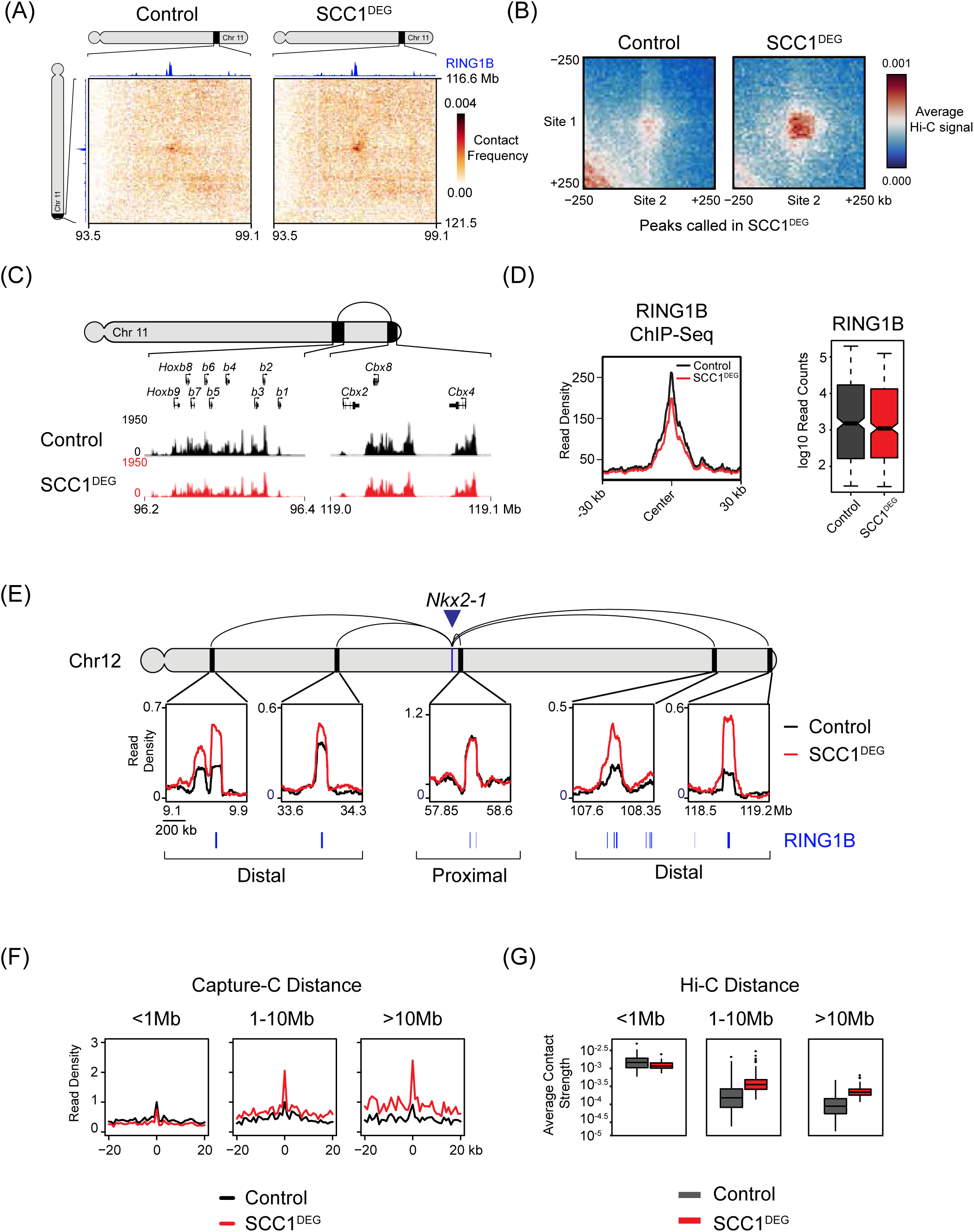
Cohesin removal strengthens long-range polycomb chromatin domain interactions. (A) Hi-C illustrating an interaction that increases in strength in the SCC1^DEG^ cell line visualised at 20 kb resolution. RING1B ChIP-seq is displayed above and to the left of the matrices. (B) Aggregate analysis of Hi-C from Control and SCC1^DEG^ cells at peaks that persist in the absence of cohesin (n=336). (C) ChIP-Seq for RING1B in the Control and SCC1^DEG^ cells at the interacting sites shown in (A). (D) RING1B ChIP-seq signal (metaplots (left) and boxplots (right)) at RING1B peaks overlapping interactions that persist in the absence of cohesin. (E) Capture-C interaction profiles between the *Nkx2-1* promoter and selected proximal and distal RING1B-occupied sites in the Control and SCC1^DEG^ cells. RING1B ChIP-seq peaks are shown as blue bars below. The location of the *Nkx2-1* promoter is indicated with a blue arrow/bar and the interactions sites as black bars on the chromosome. Read density corresponds to normalized reads in the capture averaged across 250 DpnII restriction fragments. (F) Aggregate Capture-C signal in the Control and SCC1^DEG^ cells at interaction sites segregated based on distance from the capture site. Only interactions between polycomb target gene promoters and RING1B occupied sites present in SCC1^DEG^ are shown. Read density was normalised to Control signal at the summit and the x-axis illustrates the distance from the interaction site in DpnII fragments. (G) Average Hi-C contact strength in the Control and SCC1^DEG^ at interactions that persist in the absence of cohesin segregated based on distance between the interactions.

To further explore polycomb-dependent interactions and their regulation by cohesin we used a technique called Capture-C that has an advantage over Hi-C in providing increased sensitivity and resolution for interrogating interactions at specific regions in the genome (Hughes et al., 2014). Using Capture-C we focussed on 18 genes that are associated with polycomb chromatin domains and examined their interactions following removal of cohesin. Interestingly, our analysis demonstrated that interactions between polycomb chromatin domains that are in close proximity tended to be unchanged or slightly reduced in the absence of cohesin (Figure 3E and F). In contrast, interactions that occurred over long distances, often between different TADs, showed increases in their interaction strength (Figure 3E and F). Importantly, these interactions were lost following PRC1 removal, demonstrating that they rely on intact polycomb chromatin domains (Figure S3A and B). This distance dependent effect was also evident when we examined interactions in our Hi-C analysis (Figure 3G). Therefore, cohesin has little effect on interactions between polycomb chromatin domains that are in close proximity on the chromosome, but counteracts interactions between polycomb chromatin domains separated by large distances.

### Cohesin counteracts polycomb chromatin domain interactions independently of TADs and insulation

The effect of cohesin loss on polycomb chromatin domain interactions could be related to loss of TADs or alternatively TAD-independent processes. To distinguish between these possibilities, we carried out Capture-C in a cell line where TADs are disrupted by removal of CTCF rather that removal of cohesin (Figure 4A and B and S4) (Nora et al., 2017). In contrast to the loss of cohesin, removal of CTCF did not strengthen distal polycomb chromatin domain interactions (Figure 4D and E). This was also evident when we examined the interactions that persisted in the absence of cohesin after CTCF removal using Hi-C (Nora et al., 2017) (Figure 4C and F). Therefore, cohesin counteracts polycomb chromatin domain interactions through a process that is independent of CTCF and TADs.

**Figure 4.**
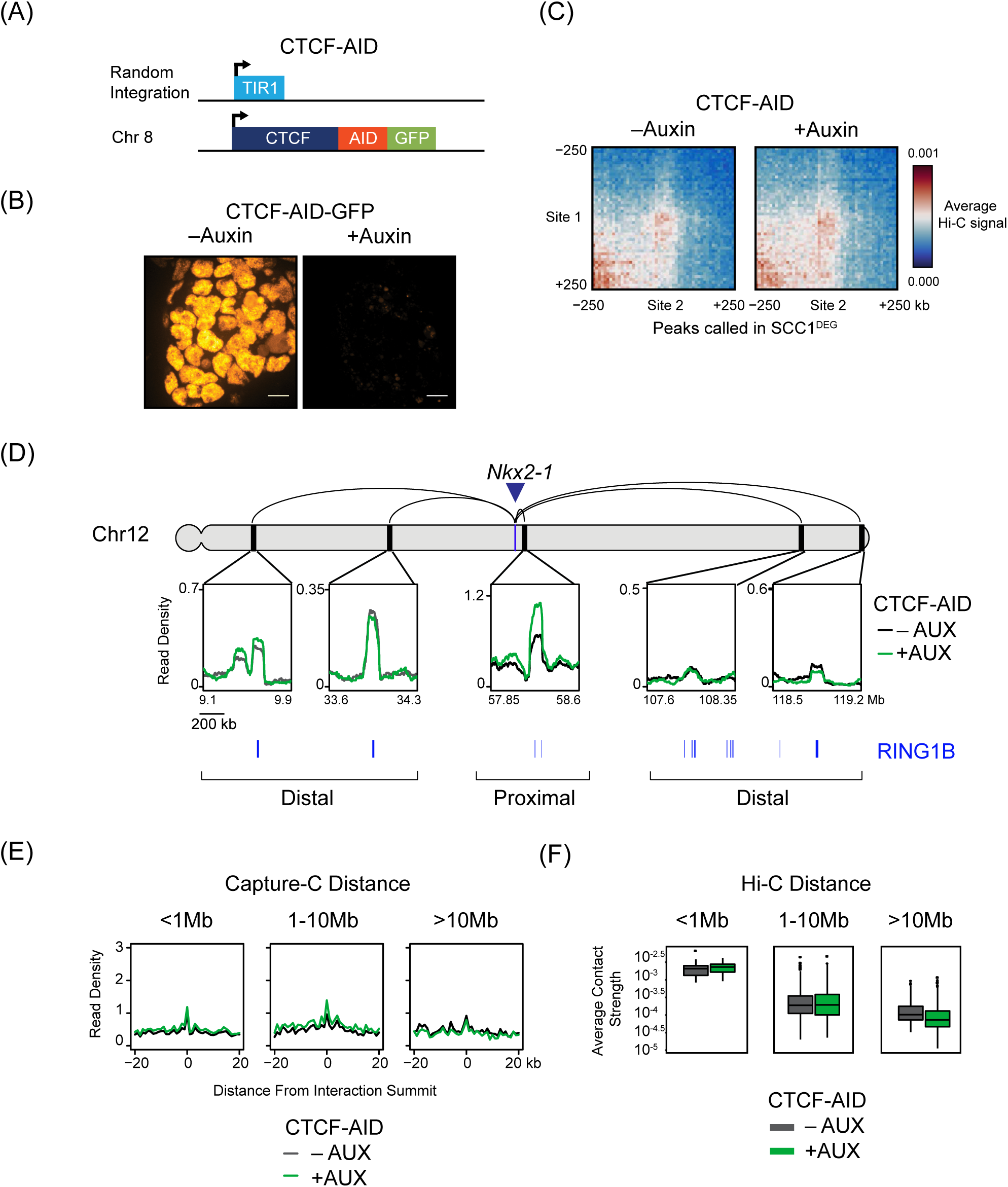
Cohesin counteracts polycomb chromatin domain interactions independently of TADs and insulation. (A) A schematic illustrating the genotype of the CTCF-AID-GFP cell line (Nora et al., 2017). (B) Live cell microscopy images of CTCF-AID-GFP cells ± auxin (48 h). Scale bar = 10 µm (bottom). (C) Aggregate analysis of Hi-C from CTCF-AID-GFP cells ± auxin (Nora et al., 2017) at interactions that persist in the absence of cohesin (n=336). (D) Capture-C interaction profiles between the *Nkx2-1* promoter and selected proximal and distal RING1B-occupied sites in the CTCF-AID-GFP cells ± auxin. RING1B ChIP-seq peaks are shown as blue bars below. The location of the *Nkx2-1* promoter is indicated with a blue arrow/bar and the interactions sites as black bars on the chromosome. Read density corresponds to normalised reads in the capture averaged across 250 DpnII restriction fragments. (E) Aggregate Capture-C signal in the CTCF-AID-GFP cells ± auxin at interactions that persist in the absence of cohesin segregated based on distance from the capture site. Only interactions between polycomb target gene promoters and RING1B occupied sites present in SCC1^DEG^ are shown. Read density was normalised to signal at the summit in CTCF-AID-GFP cells without auxin and the x-axis illustrates the distance from the interaction site in DpnII fragments. (F) Average Hi-C contact strength in the CTCF-AID-GFP cells ± auxin at interactions that persist in the absence of cohesin segregated based on distance between the interactions.

### Polycomb chromatin domain interactions are disrupted by cohesin

Chromosome conformation capture-based approaches are extremely sensitive and can identify infrequent interaction events like those that lead to the emergence of TADs in ensemble Hi-C analysis. However, neither Hi-C nor Capture-C reveal the absolute frequency of these interactions. Therefore, to characterise polycomb chromatin domain interactions and define the extent to which cohesin regulates these, we set out to measure interactions in single cells. We focused on a pair of genes (*HoxD10* and *Dlx2*) with polycomb chromatin domains that showed increased interaction in Hi-C (Figure 1C) and Capture-C (Figure 5A) after cohesin removal. We generated probes that uniquely mark the *HoxD10* and *Dlx2* genes and performed non-denaturing RASER-FISH (Brown et al., 2018) to measure the three-dimensional distance between these loci in individual cells (Figure 5B and S5C). This revealed a distribution of distances between *HoxD10* and *Dlx2* in cells where cohesin is intact, including some in close proximity (Figure 5C and S5D). When cohesin was removed, the number of very close distances became larger, in agreement with increased contact probabilities between polycomb chromatin domains observed in Hi-C and Capture-C (Figure 3). Importantly, to test whether this was dependent on both the inactivation of cohesin and the presence of PRC1, we developed a double degron line where SCC1 and RING1B were degraded simultaneously by addition of auxin (Figure S5A). Capture-C in this line revealed a loss of interaction between *HoxD10* and *Dlx2* (Figure S5B). Similarly, in the absence of both cohesin and PRC1 the number of very close distances between *HoxD10* and *Dlx2* in FISH was greatly reduced (Figure 5C). Importantly, similar effects were observed when we examined the interaction of polycomb chromatin domains associated with the *Nkx2-3* and *Pax2* genes (Figure S6). Together this demonstrates that cohesin counteracts polycomb chromatin domain association in single cells.

**Figure 5.**
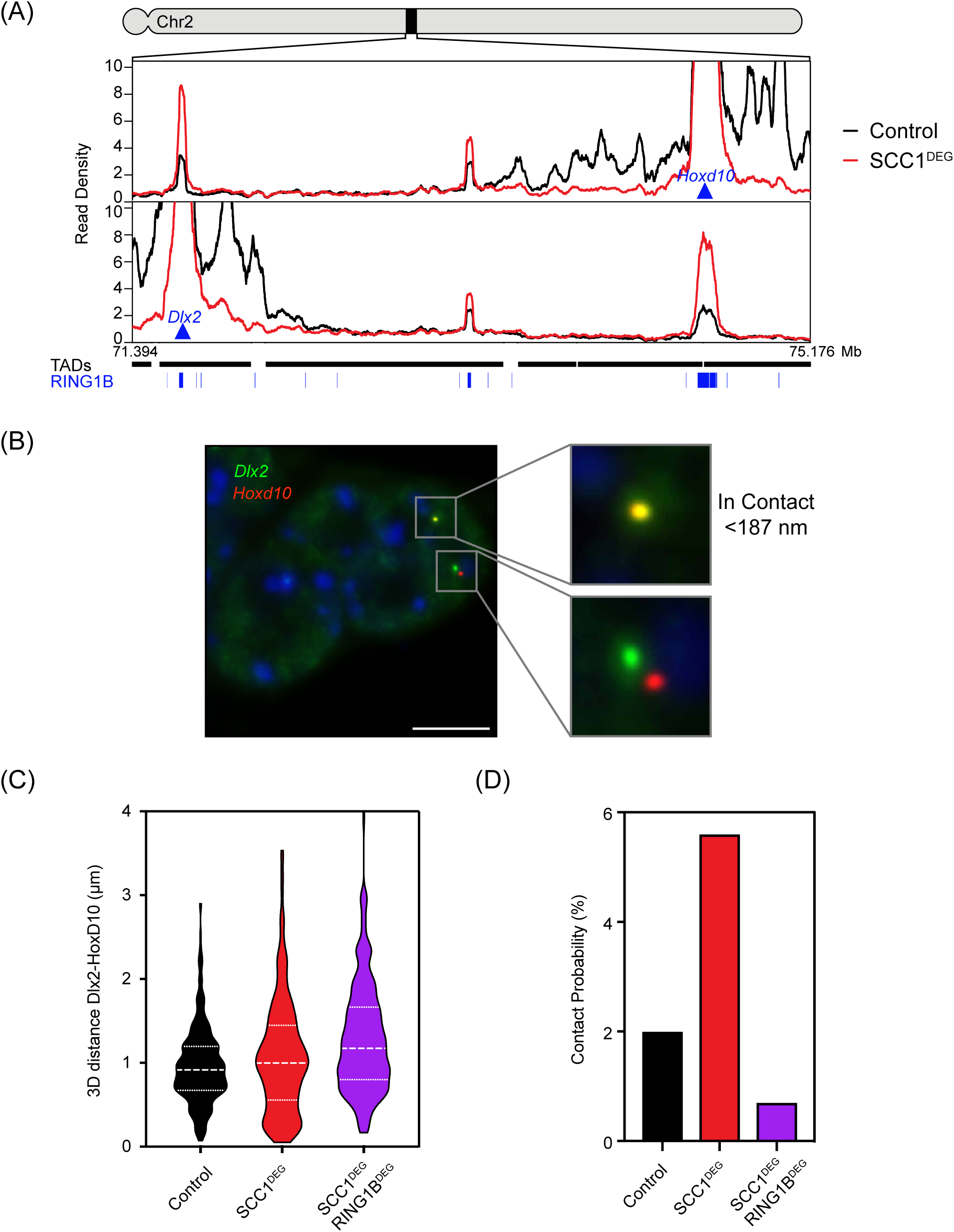
Polycomb chromatin domain interactions are disrupted by cohesin. (A) Capture-C interaction profiles from *Hoxd10* (top) and *Dlx2* (bottom) viewpoints in Control and SCC1^DEG^ lines. RING1B ChIP-Seq peaks are displayed as blue bars and TAD intervals as black bars. (B) Representative image of RASER-FISH showing signals classed as in contact (top pair 0.0905 µm apart) and not in contact (bottom pair 1.1629 µm apart). Probes are for *Dlx2* (green) and *HoxD10* (red). Scale bar = 5 µm. (C) Violin plots showing 3D distance measurements between *Dlx2* and *HoxD10* in the indicated cells lines. The dashed lines show the median and interquartile range of between n=376-409 cells for each cell line. (D) Absolute contact probabilities showing the percent of signals judged as colocalised from observations in (B) (see Methods).

Previous studies have reported that loci which display strong interactions in Hi-C do not always equate to frequent interactions in single cells (Bonev et al., 2017; Fudenberg and Imakaev, 2017). Therefore we wanted to accurately quantitate the frequency of polycomb chromatin domain interactions by determining the number of FISH probe measures that may be considered to be in contact (Cattoni et al., 2017). This revealed that *HoxD10* and *Dlx2* polycomb chromatin domains were in contact (closer than 187nm) 2% of the time (Figure 5D). Following cohesin removal, the contact frequency increased to 5.6% indicating that when cohesin is present it functions to disrupt interactions between regions of chromatin occupied by the polycomb repressive system. Importantly, this increased association between polycomb chromatin domains was dependent on cohesin and PRC1, as their simultaneous removal reduced the interaction frequency to 0.7%. Again, we observed very similar interaction frequencies when we examined the polycomb chromatin domains associated with the *Nkx2-3* and *Pax2* genes (Figure S6C). It is also important to point out that these interaction values likely underestimate the absolute frequency with which polycomb chromatin domains interact with one another, as interactions between specific pairs of sites are likely to vary between individual cells. Although this potential underestimation is not accounted for in FISH analysis, it is consistent with imaging of polycomb proteins in ESCs where hundreds of cytologically distinct foci that house polycomb occupied genes, called polycomb bodies are evident (Isono et al., 2013). Together these single cell measurements quantitate the frequency with which polycomb chromatin domains interact in single cells and demonstrate that cohesin disrupts polycomb chromatin domain interactions.

### Increased polycomb chromatin domain association in the absence of cohesin supresses gene expression

In vertebrates, polycomb repressive complexes play important roles in maintaining the repression of genes in cell types where they should not be expressed (Schuettengruber et al., 2017). This is proposed to rely on chromatin modifications and, in some instances, on the formation of polycomb-dependent chromatin interactions (Eskeland et al., 2010; Kundu et al., 2017). Here we demonstrate that cohesin counteracts and disrupts long-range interactions between polycomb chromatin domains and their associated genes. We were therefore interested to test whether cohesin affects polycomb-mediated gene repression. To examine this possibility we performed calibrated RNA-Seq (cRNA-seq) before and after cohesin removal. In agreement with previous analysis following cohesin depletion, changes in gene expression were modest and the transcription of only several hundred genes was significantly altered (Rao et al., 2017) (Figure 6A). Nevertheless, we also observed a more subtle and widespread reduction in gene transcription in agreement with a proposed role for cohesin in supporting promoter-enhancer interactions and gene expression (Hadjur et al., 2009; Schwarzer et al., 2017). Remarkably, however, RING1B-bound genes were overrepresented (251/365) among the genes whose expression was significantly reduced, indicating that they were disproportionally affected (Figure 6B). A more detailed analysis of polycomb bound genes with detectable expression (Figure S7A) in our cRNA-seq showed that reductions in expression were larger in magnitude following cohesin removal if the gene interacted with another polycomb chromatin domain (Figure 6C and D). Together these observations reveal that cohesin, and presumably its loop extruding activity, play a direct role in counteracting long-range polycomb chromatin domain interactions and gene repression.

**Figure 6.**
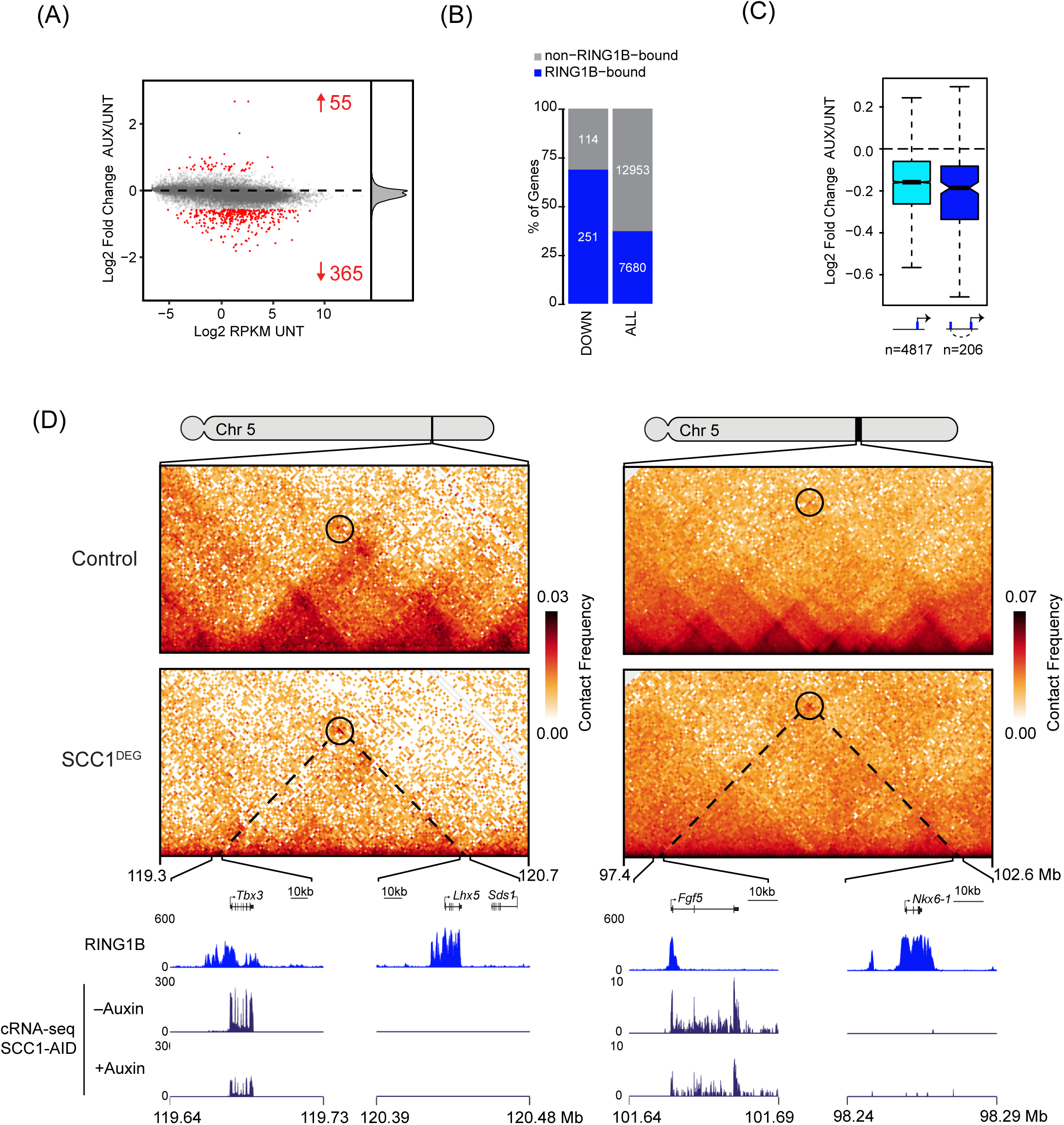
Increased polycomb chromatin domain association in the absence of cohesin suppresses gene expression. (A) An MA plot of gene expression alterations in Scc1-mAID-GFP cells ± auxin (6 h). The number of genes with increased or decreased expression (p-adj < 0.05 and > 1.5-fold) is shown in red. The density of Log2 Fold changes is shown on the right. (B) RING1B binding (+/-1kb from the TSS, blue bars) at gene promoters that show reductions in gene expression following cohesin removal (left) compared to all genes (right). Empirical p-value for RING1B-bound genes enrichment within the downregulated genes: p=0 (n=10000 random tests). (C) The magnitude of gene expression change at expressed RING1B bound genes that do (right) or do not (left) interact with another RING1B bound site in Hi-C. (D) Hi-C (left at 40 kb and right at 10 kb resolution), cRNA-Seq and RING1B ChIP-Seq for two examples of genes with interactions in Hi-C that are strengthened after cohesin removal and whose gene expression decreases.

## Discussion

How cohesin functions to shape chromosome structure and function remains poorly understood. Here, using degron alleles and chromosome conformation capture approaches, we identify a series of long-range interactions that persist in the absence of cohesin and correspond to polycomb chromatin domains (Figure 1). We demonstrate that PRC1 is essential for the formation of these interactions (Figure 2). Remarkably, in the absence of cohesin, polycomb chromatin domain interactions are strengthened, revealing that they are normally counteracted by cohesin (Figure 3). Importantly, cohesin regulates these interactions independently of CTCF and TADs (Figure 4). Using cellular imaging we visualise polycomb chromatin domain interactions, quantify their frequency, and further demonstrate a role for cohesin in separating polycomb chromatin domains and regulating their interaction in single cells (Figure 5). Finally, we demonstrate that regulation of polycomb chromatin domain interactions by cohesin affects gene expression (Figure 6). These findings reveal a new link between the capacity of the polycomb system to form long-range transcriptionally repressive chromosome interactions, and cohesin which appears to actively counteract and regulate this process (Figure 7).

**Figure 7.**
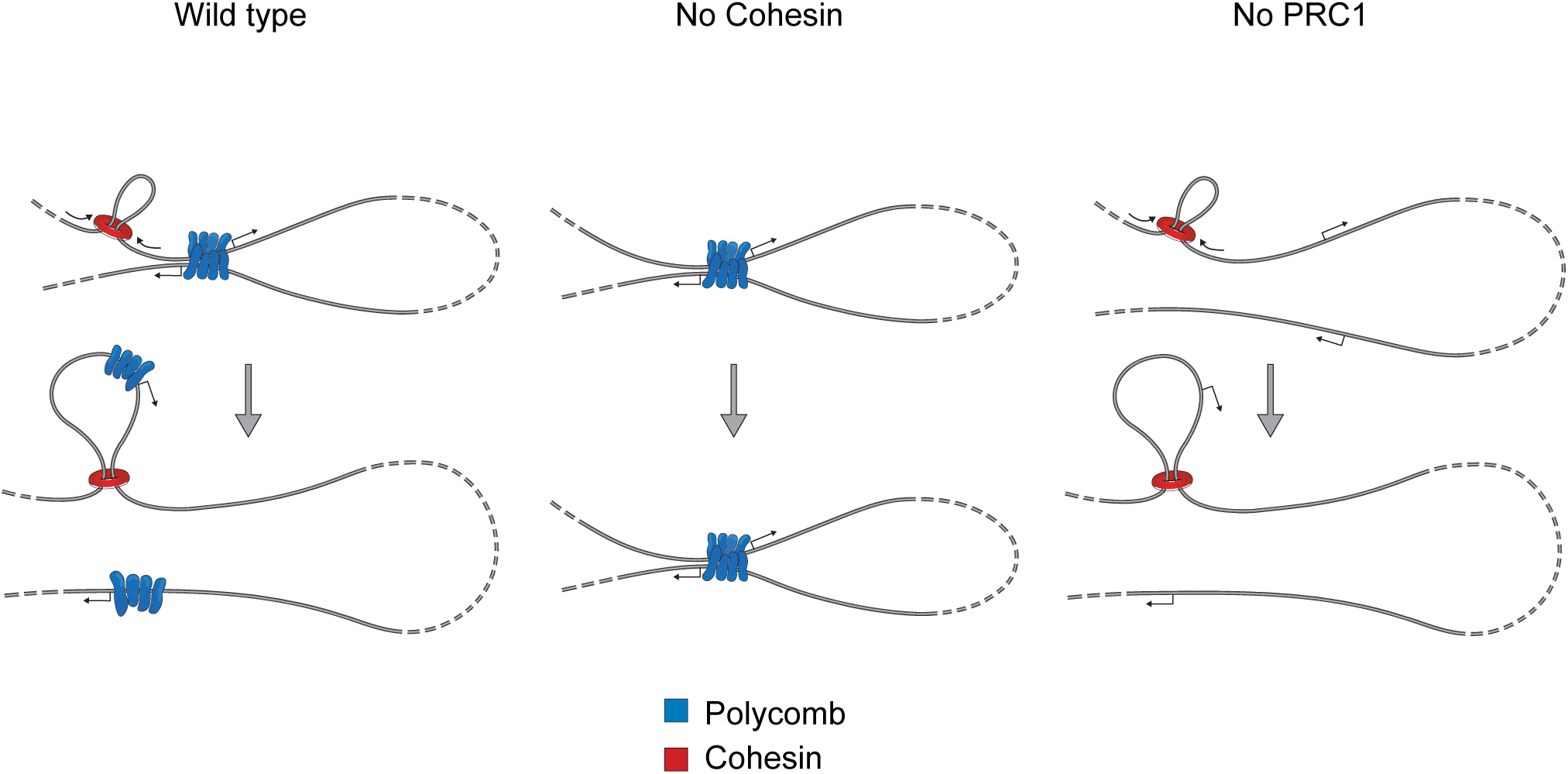
A model for disruption of polycomb chromatin domain interactions by cohesin and loop extrusion. Polycomb chromatin domains (blue) can interact even when separated by large distances on the chromosome. Cohesin (red circle) can load onto DNA and has been proposed to extrude chromatin. As loop extrusion proceeds it will encounter one of the two interacting polycomb chromatin domains. We propose that the manner in which chromatin is extruded through cohesin could lead to the individualisation of these two previously interacting polycomb chromatin domains and explain the observed effect that cohesin removal has on polycomb chromatin domain interaction in our chromosome conformation capture and single-cell imaging experiments. A key prediction of this model would be that cohesin loading and extrusion near either interacting polycomb chromatin domain would lead to the observed effect.

Initially these observations may seem counterintuitive. Why would it be advantageous for a cell to disrupt chromosomal interactions, like those formed between polycomb chromatin domains, which function to protect against inappropriate gene expression? One simple explanation may be that, if left unchecked, progressive association and possibly compartmentalisation by factors which nucleate and promote repressive chromatin interactions could lead to an irreversibly silent state. This may be particularly pertinent in the case of the polycomb system, as recently components of PRC1 have been shown to phase separate, and this has been linked to selective exclusion of gene regulatory factors (Plys et al., 2018; Tatavosian et al., 2019). In pluripotent cells, or at early developmental stages, such a static situation could be deleterious, as many genes occupied by polycomb chromatin domains and which engage in long-range interactions must be expressed later in development. It is tempting to speculate that cohesin primarily functions on interphase chromosomes to counteract the potential for such stasis by periodically breaking up self-associating structures and in doing so provide an opportunity for factors in the nucleus to constantly sample these regions of the genome should they be required for future gene expression programmes.

Interestingly, interactions between super-enhancers in cancer cells were previously shown to occur independently of cohesin and these elements also appear to have a tendency to phase separate (Sabari et al., 2018). Similar to the increased association we observe between polycomb chromatin domains, long-range super-enhancer associations were also increased following cohesin removal (Rao et al., 2017). Conceptually aligned with the idea that cohesin and loop extrusion may counteract polycomb chromatin domain interactions to mitigate stasis, one could envisage how periodically disrupting super-enhancer associations and possibly their interactions with gene promoters might support a constant re-evaluation of gene regulatory interactions. Again, this could provide an opportunity for plasticity in transitioning between gene expression programmes.

As chromosomal interactions are studied in more detail it is becoming evident that TADs do not correspond to fixed or invariant structures in single cells. Instead, they appear to emerge in ensemble Hi-C analysis from low frequency tendencies to interact across many cells. Spatially heterogeneous globular chromatin structures, similar in size to TADs, are evident in single cells, but form independently of cohesin. This indicates that cohesin and loop extrusion do not primarily function to create structure in chromosomes. In agreement with these observations, our results would argue that cohesin and loop extrusion instead disrupt chromatin interactions through constantly separating regions of chromatin and increasing chromosomal dynamics. This may rely on the topological manner in which entrapped chromatin would extrude through the cohesin complex, however translocation of cohesin through polycomb domains by other mechanisms could also disrupt interactions. We envisage that loading of cohesin on the chromosome in proximity to a polycomb chromatin domain, followed by loop extrusion, could break up interactions with other polycomb chromatin domains, irrespective of whether they are separated by large distance on the chromosome or even between chromosomes. This would also explain why CTCF and its proposed activity in halting extrusion would not affect the ability of cohesin to counteract polycomb chromatin domain interactions as we observe. Instead, CTCF and termination of loop extrusion, may function to restrict the activity of gene regulatory elements to regions between CTCF sites by limiting mixing of chromatin that might result from unconstrained loop extrusion.

Finally, cohesin is best characterised for the role it plays in holding sister chromatids together after replication and during cell division. In contrast, other SMC complexes, for example bacterial SMC-ScpAB and eukaryotic condensin, have been proposed to play roles in separating chromosomes through processes that are thought to rely on loop extrusion (Goloborodko et al., 2016; Nasmyth, 2001; Wang et al., 2017). Our observations provide new evidence to suggest that in addition to its role in sister chromatid cohesion, cohesin also retains its primordial SMC complex activity in separating regions of chromosomes as is evident from the role it plays in disrupting long-range polycomb chromatin domain interactions.

## Acknowledgements

We would like to thank Elphège Nora and Benoit Bruneau for providing the CTCF degron cell line. We would like to thank Neil Brockdorff and Francis Barr for providing comments on the manuscript. We are grateful to Amanda Williams at the Department of Zoology, Oxford, for sequencing support on the NextSeq 500. We are grateful to Jim Hughes, Damien Downes and Priscila Hirschfeld for providing advice, protocols and probes for Capture-C prior to publication.

## Figure Legends

**Figure S1.**
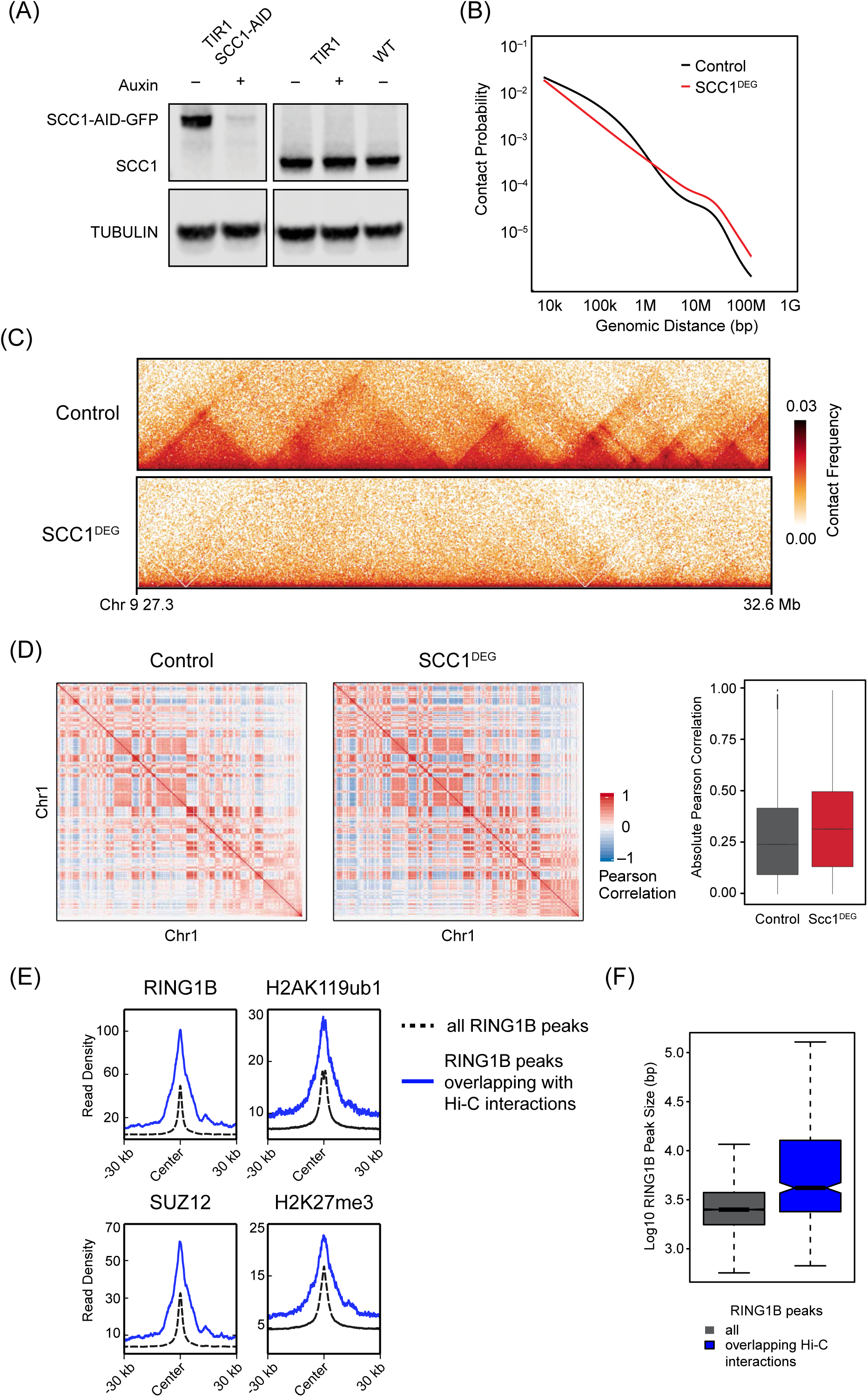
(A) A representative western blot for SCC1 in the TIR1 and SCC1-mAID-GFP (=SCC1^DEG^) cell lines ± auxin. A wild type cell line is shown for comparison and tubulin is shown as a loading control. (B) Genomic distance-dependent contact probability from Hi-C in Control or SCC1^DEG^ cells. (C) Hi-C in Control and SCC1^DEG^ cells at 10 kb resolution. (D) Pearson correlation coefficient of chromosome 1 from Control and SCC1^DEG^ at 500 kb resolution (left). Bar plot of the genome-wide absolute Pearson correlation for Control and SCC1^DEG^ (right). (E) RING1B, H2AK119ub1, SUZ12, and H3K27me3 ChIP-seq signal metaplotted at RING1B peaks overlapping with interactions that persist in the absence of cohesin (blue line) or all RING1B peaks (dashed-black line). (F) A box plot showing the RING1B peak size at RING1B peaks overlapping interactions that persist in the absence of cohesin (blue right) or all RING1B peaks (grey left).

**Figure S2.**
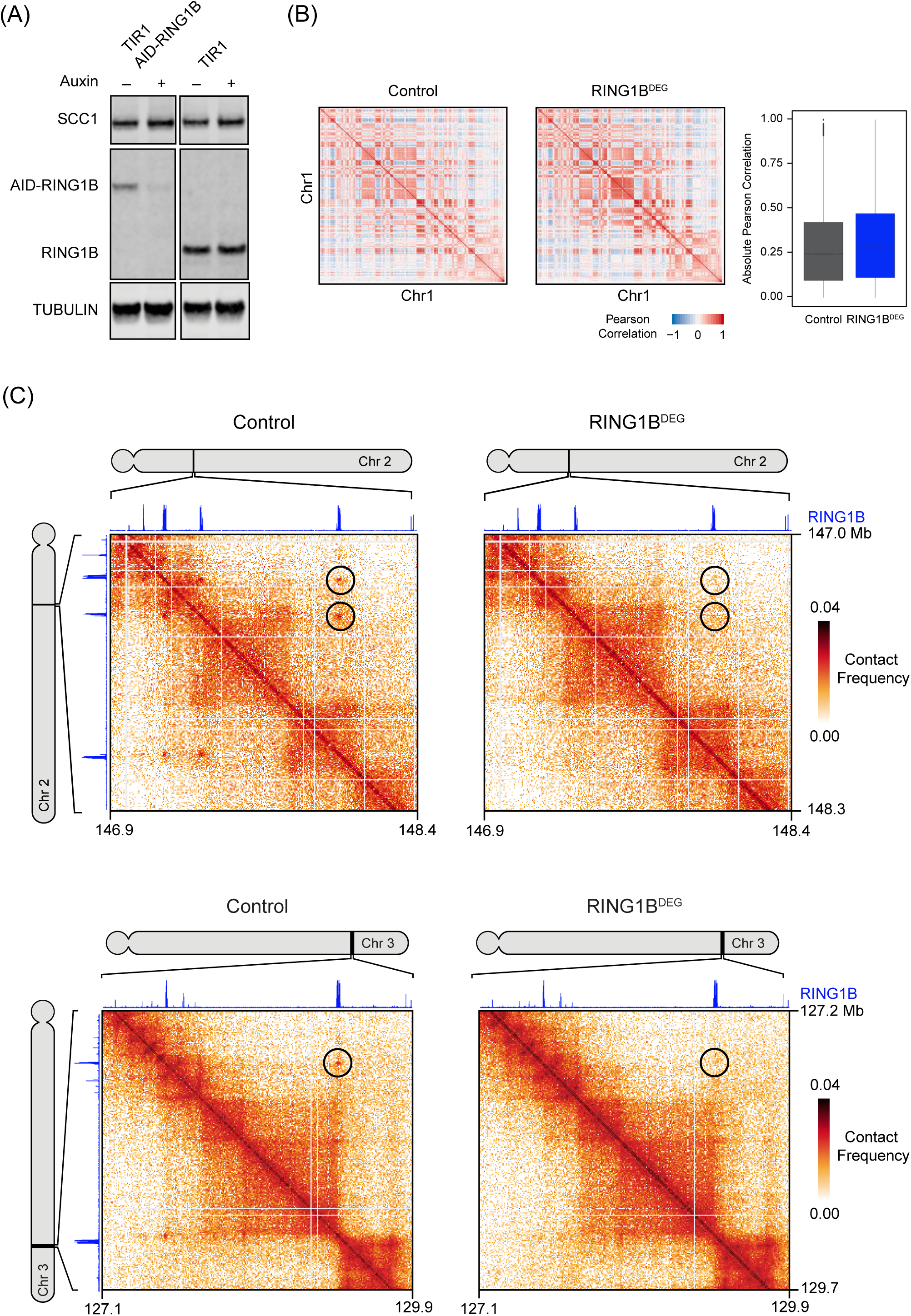
(A) A representative western blot for SCC1 and RING1B in the Control and AID-RING1B cell lines ± auxin. Tubulin is shown as a loading control. (B) Pearson correlation coefficient of chromosome 1 from Control and RING1B^DEG^ at 500 kb resolution (left). Bar plot of the genome-wide absolute Pearson correlation for Control and RING1B^DEG^ (right). (C) Hi-C in Control and RING1B^DEG^ cells at two regions (top at 5 kb resolution and bottom at 10 kb resolution). Black circles indicate at interactions that persist in the absence of cohesin and RING1B ChIP-seq is displayed above and to the left of the matrices.

**Figure S3.**
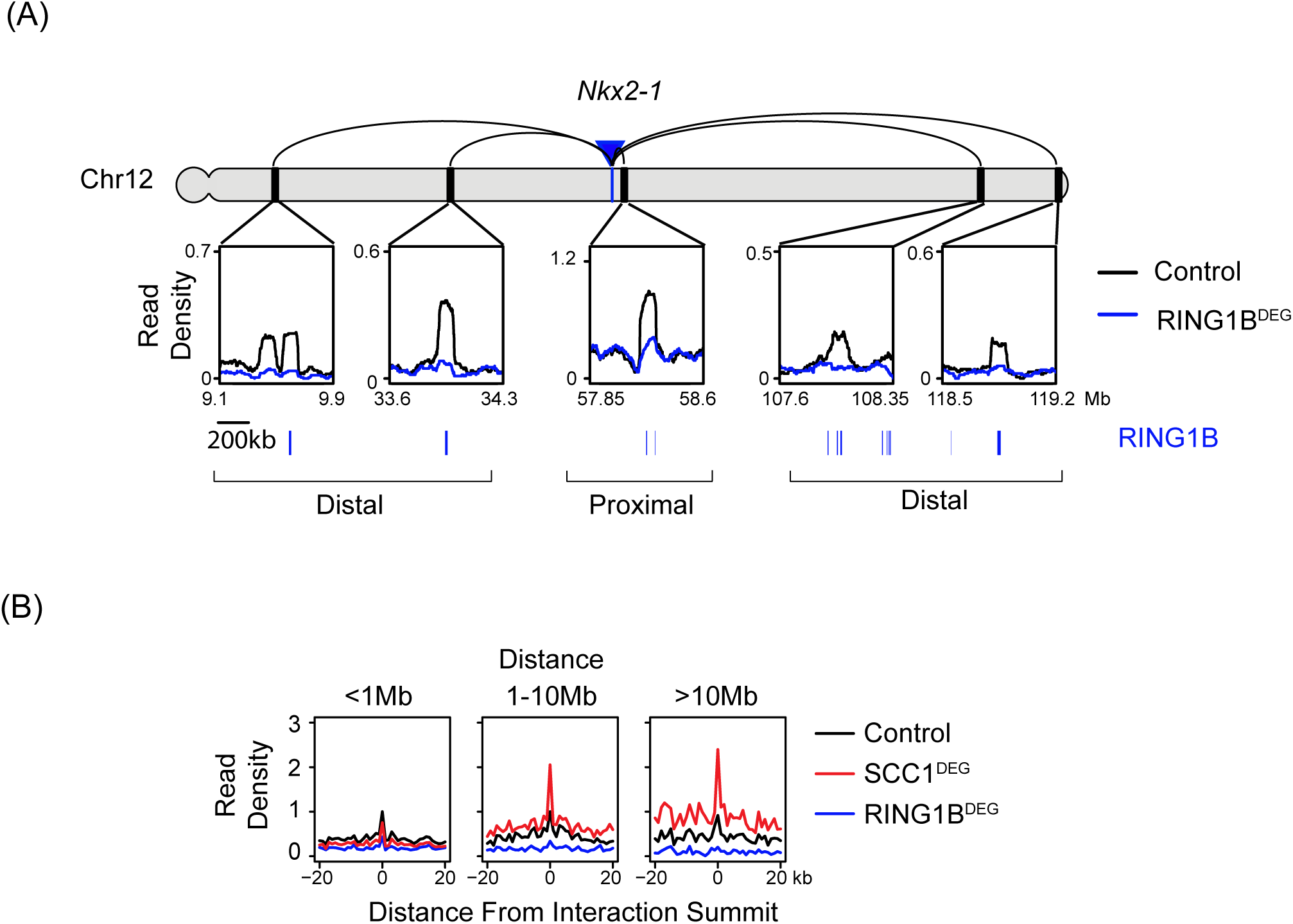
(A) Capture-C interaction profiles between the *Nkx2-1* promoter and selected proximal and distal RING1B-occupied sites in the Control and RING1B^DEG^ cell lines. RING1B ChIP-seq peaks are shown as blue bars below. The location of the *Nkx2-1* promoter is indicated with a blue arrow/bar and the interactions sites as black bars on the chromosome. Read density corresponds to normalised reads in the capture averaged across 250 DpnII restriction fragments. (B) Aggregate Capture-C signal in the Control, SCC1^DEG^ and RING1B^DEG^ cells at interaction sites segregated based on distance from the capture site. Only interactions between polycomb target gene promoters and RING1B occupied sites present in SCC1^DEG^ are shown. Read density was normalised to Control signal at the summit and the x-axis illustrates the distance from the interaction site in DpnII fragments.

**Figure S4.**
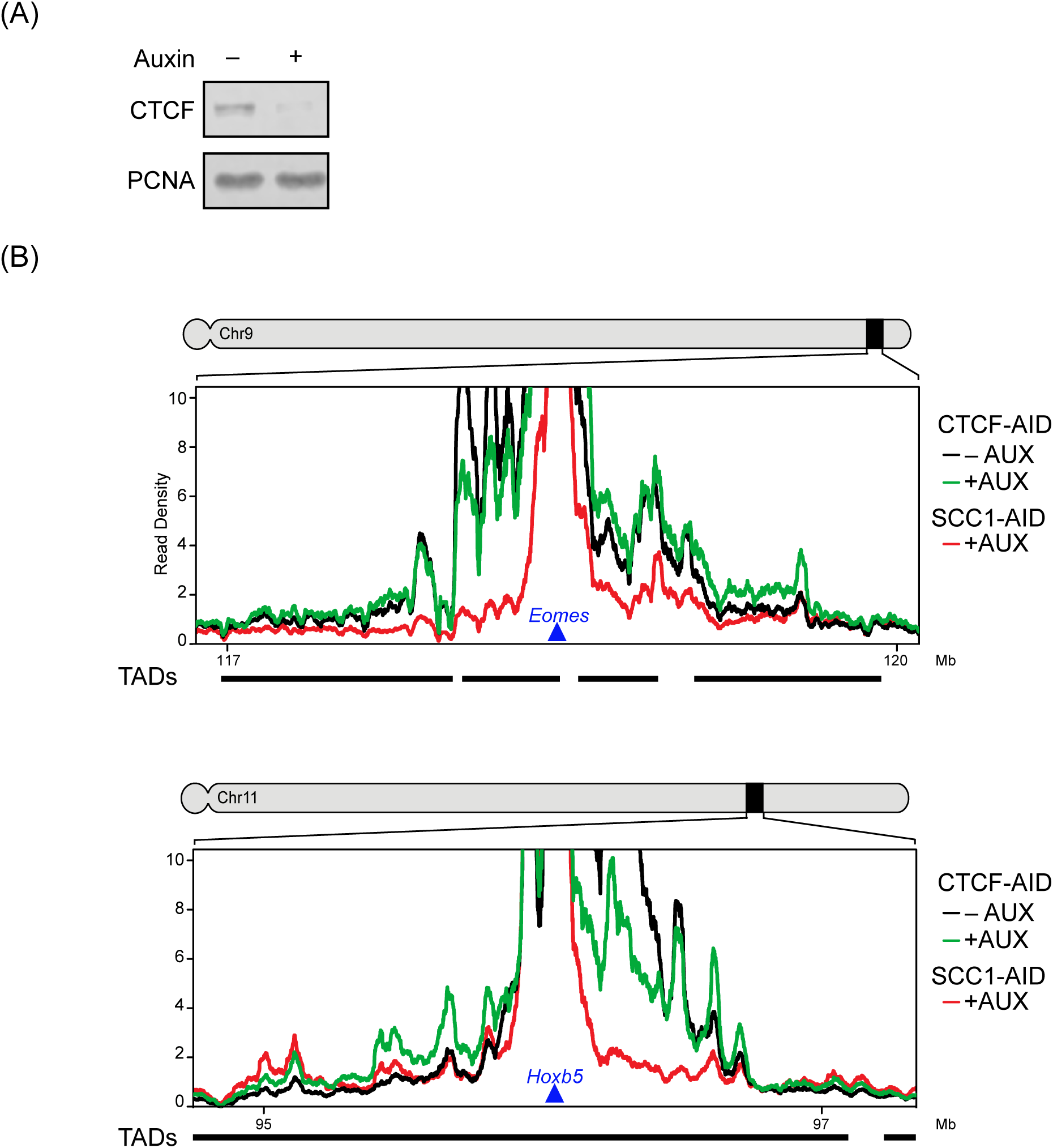
(A) A representative western blot for CTCF in the CTCF-AID cell lines ± auxin. PCNA is shown as a loading control. (B) Capture-C signal around *Eomes* and *Hoxb5* gene promoters. Shown are normalized read densities for CTCF-AID +/-AUX (green and black, respectively) and as comparison for SCC1^DEG^ (red). Read density corresponds to normalised reads in the capture averaged across 80 DpnII restriction fragments. View point is indicated as a blue triangle. TAD boundaries are shown below.

**Figure S5.**
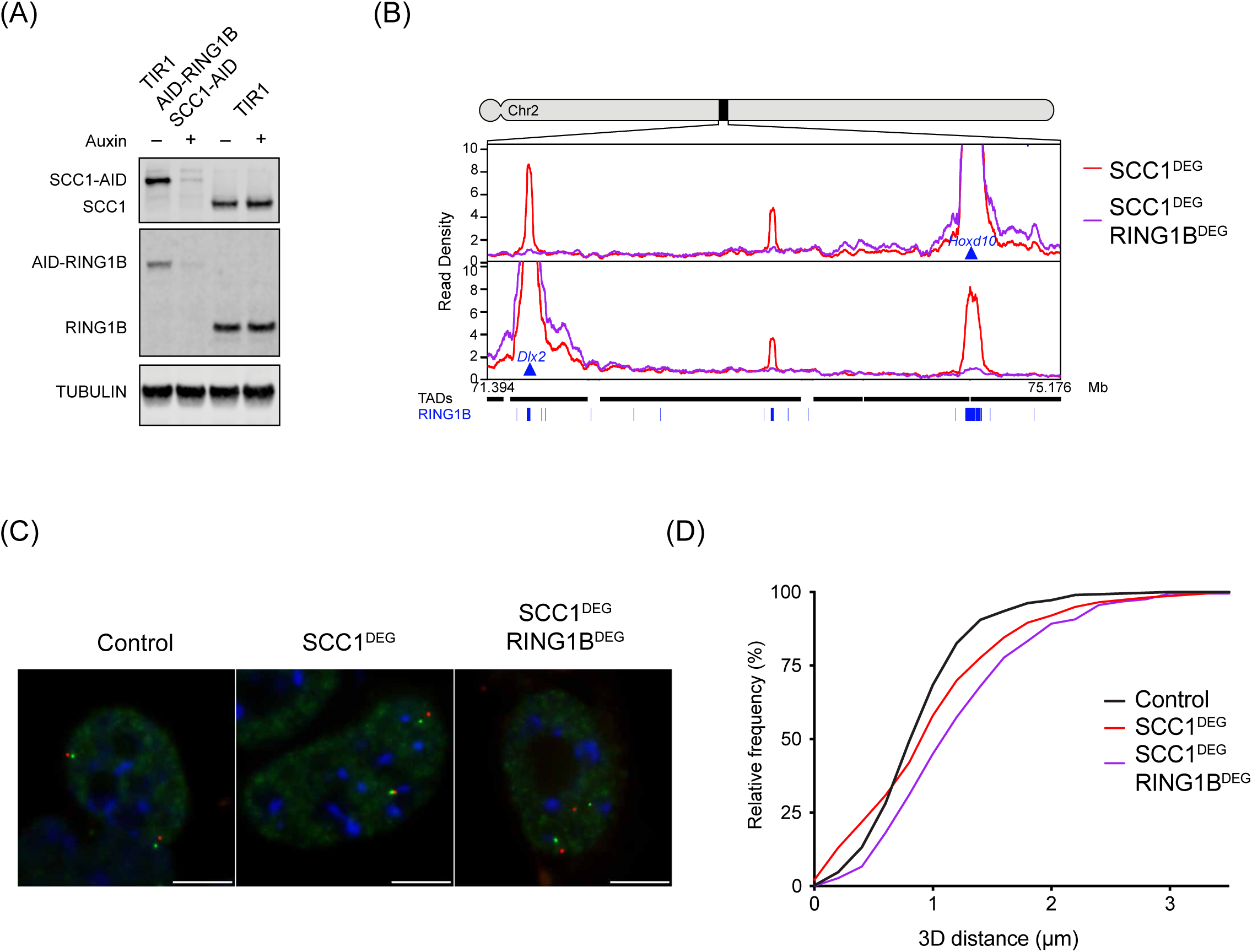
(A) A representative western blot for SCC1 and RING1B in the TIR1 and AID-RING1B SCC1-mAID-GFP cell lines ± auxin. Tubulin is shown as a loading control. (B) Capture-C interaction profiles from *Hoxd10* (top) and *Dlx2* (bottom) viewpoints in Control and SCC1^DEG^ RING1B^DEG^ cell lines. RING1B ChIP-Seq peaks are displayed as blue bars and TAD intervals are as black bars. (C) Representative *Hoxd10* (red) *Dlx2* (green) RASER-FISH images from the indicated cell lines. Scale bar = 5 µm. (D) Cumulative frequency distribution of 3D distance measures between *Hoxd10* and *Dlx2* in the indicated cells lines. Measurements as in Figure 5C.

**Figure S6.**
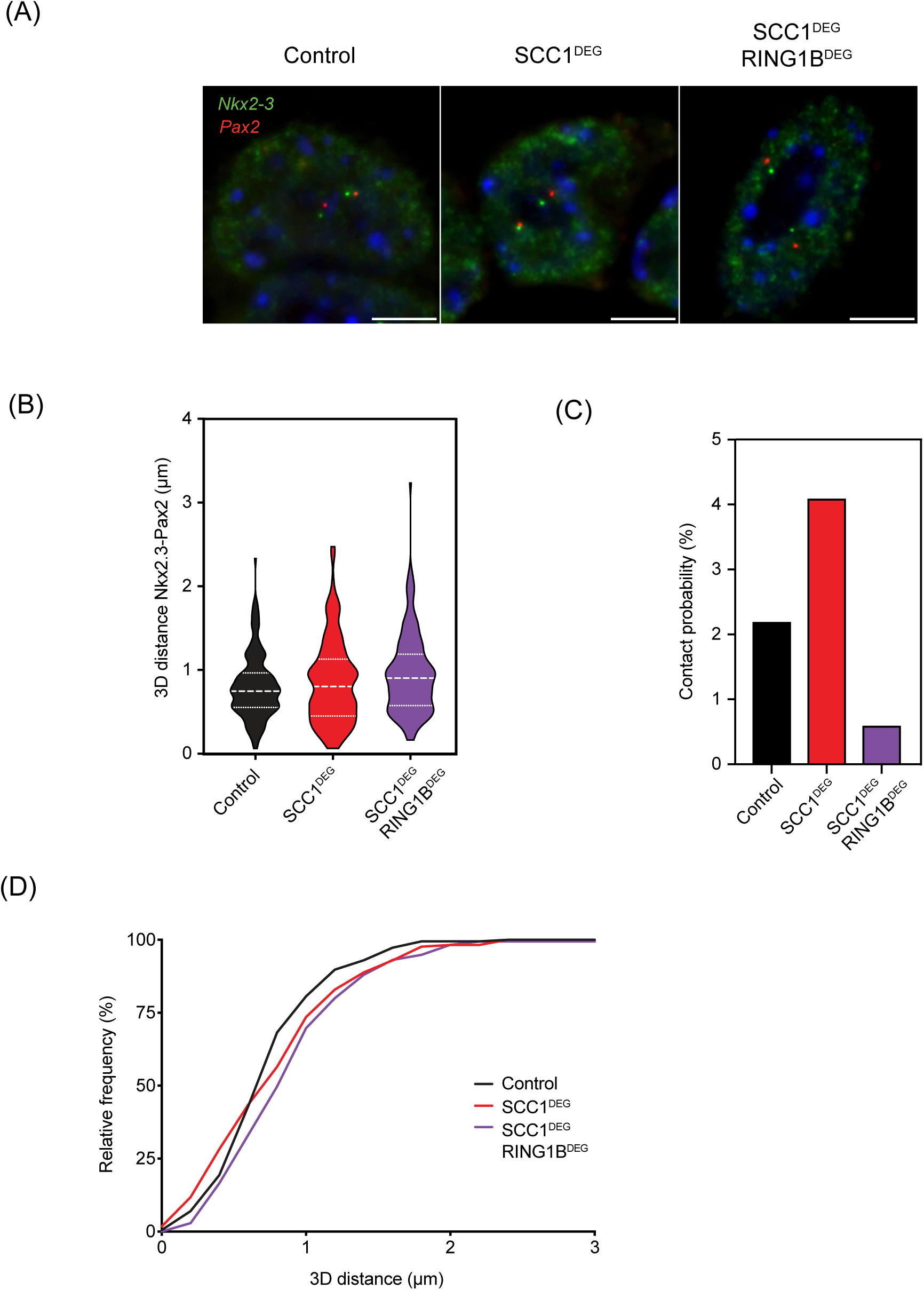
(A) Representative RASER-FISH images illustrating the *Nkx2-3* (green) and *Pax2* (red) loci. Scale bar is 5 µm. (B) Violin plots showing 3D distance measurements between *Nkx2-3* and *Pax2* in the indicated cells lines. The dashed lines show the median and interquartile range of between n=170-186 cells for each cell line. (C) Absolute contact probabilities showing the percent of signals judged as colocalised from observations in (B) (see Methods). (D) Cumulative frequency distribution of 3D distance measures between *Nkx2-3* and *Pax2* in the indicated cells lines. Measurements as in (B).

**Figure S7.**
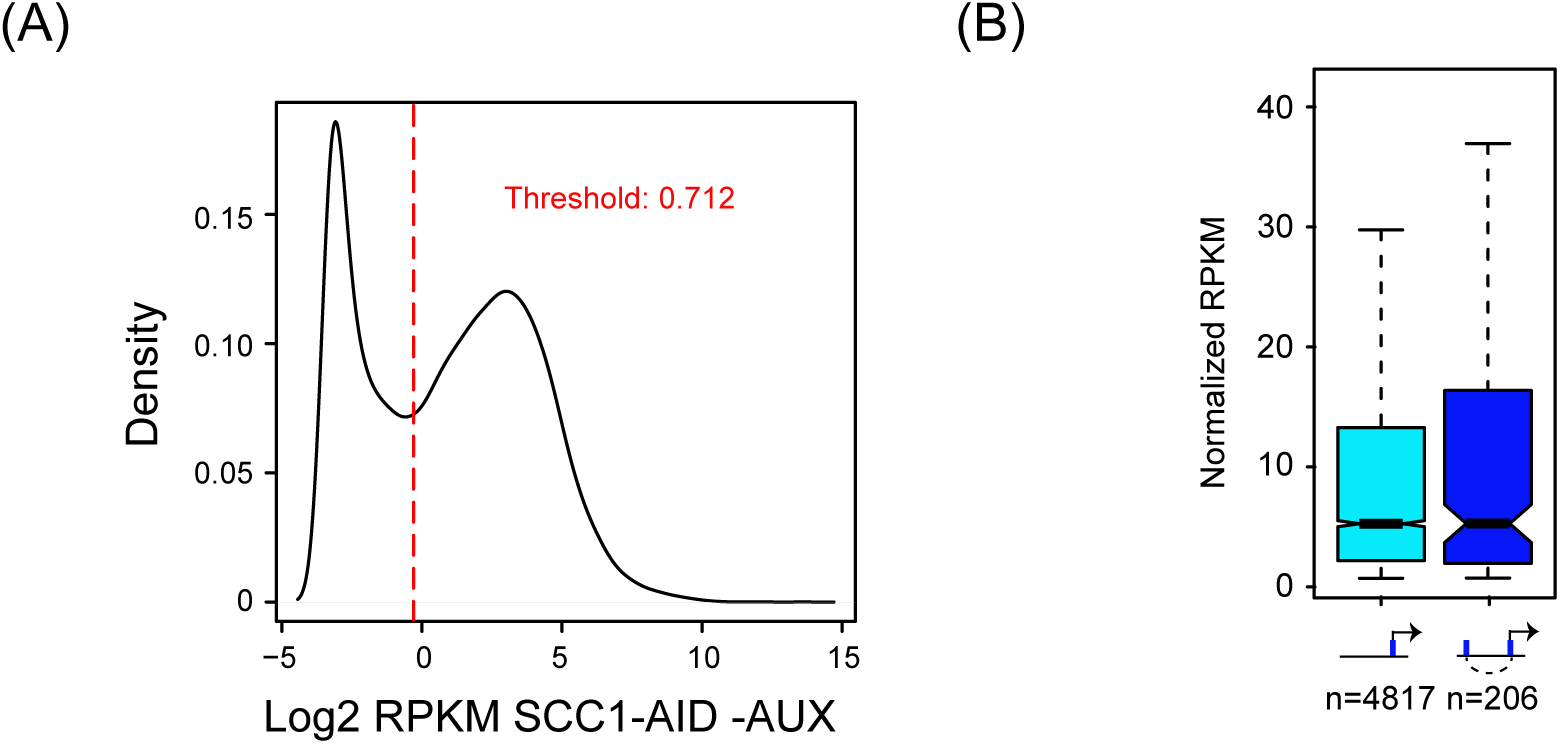
(A) Density distribution of Log2 transformed RPKM of transcripts in untreated Scc1-mAID cells. Note the bimodal distribution. Red line marks the threshold (RPKM=0.712) to distinguish expressed from unexpressed genes. (B) Boxplots show transcriptional levels in untreated Scc1-mAID cells. Transcription is shown for expressed genes with RING1B-ocupied promoters that either have no detectable Hi-C interactions in SCC1-AID cells treated with auxin (light blue), or interact with other sites that are RING1B occupied (dark blue). Numbers of genes in each group are indicated below the x-axis.

## Methods

### Cell Culture

Wild type E14 mouse ESCs cells were grown on gelatin-coated plates in DMEM supplemented with 10% FBS, penicillin/streptomycin, 5 µM 2-Mercaptoethanol, 2 mM L-glutamine, non-essential amino-acids and 10 ng/mL recombinant Leukaemia-Inhibitory Factor (LIF).

### Cloning

pSpCas9(BB)-2A-Puro (PX459) was used to construct CRISPR/Cas9 vectors (Addgene 48139). The following gRNA oligos were cloned into the BbsI restriction site:

ROSA26 CGCCCATCTTCTAGAAAGAC

SCC1 3’ CCACGGTTCCATATTATCTG

RING1A 5’UTR CTCAGCGGAGCCCCGCTTGG

RING1A Intron 3 GCGACCGTGCAGCTGACGTT

RING1B 5’ GCACAGCCTGAGACATTTCT

For homology directed gene targeting and repair 500-1000 bp homology arms were generated by Gibson Assembly.

### Gene Editing

The coding sequence for *Oryza sativa* TIR1 and a splice acceptor was heterozygously introduced into the ROSA26 locus by cotransfection of pX459 ROSA26 and pUC19 ROSA-TIR1. The resulting ESCs were ROSA-TIR1.

The mini-AID and eGFP was introduced at the C-terminus of SCC1 in ROSA-TIR1 ESCs by cotransfection of pX459 SCC1 3’ and pUC19 SCC1-mAID-eGFP. The resulting ESCs were ROSA-TIR1 SCC1-mAID-eGFP (SCC1-AID).

RING1A was deleted from ROSA-TIR1 ESCs by cotransfection of two gRNAs spanning exons 1 to 3 (pX459 RING1A 5’ UTR and pX459 RING1A Intron 3). The resulting ESCs were ROSA-TIR1 RING1AΔ.

Full length AID was introduced at the N-terminus of RING1B in ROSA-TIR1 RING1AΔ ESCs by cotransfection of pX459 RING1B 5’ and pUC19 AID-RING1B. The resulting ESCs were ROSA-TIR1 RING1AΔ AID-RING1B (RING1B-AID).

The mini-AID and GFP was introduced at the C-terminus of SCC1 in ROSA-TIR1 RING1AΔ AID-RING1B ESCs by cotransfection of pX459 SCC1 3’ and pUC19 SCC1-mAID-GFP. The resulting ESCs were ROSA-TIR1 RING1AΔ AID-RING1B SCC1-mAID-GFP (SCC1-AID RING1B-AID).

Cells were transfected using lipofectamine 2000. The next day cells were passaged and transfected cells were selected with puromycin (1 µg/ml) for two days. Eight days after puromycin removal, colonies were picked and genotyped by PCR and western blotting.

### Protein Degradation for Hi-C, Capture-C, RNA-seq and ChIP-Seq

ESCs were plated on 10 cm dishes one day before treatment. Medium was replaced with equilibrated (37°C and 5% CO_2_) medium containing auxin sodium salt (500 µM) (Sigma). The cells were incubated for 6 h or 48h (CTCF-AID) before trypsinisation and cell counting (in auxin containing medium). 100,000 cells were used for Hi-C and the rest for western blotting to confirm protein degradation.

### Hi-C

#### *In situ* Hi-C library generation for low cell input

We performed *in situ* Hi-C on control (TIR1+Auxin), SCC1^DEG^ (SCC1-AID+Auxin), RING1B^DEG^ (RING1B-AID+Auxin) ESCs (Díaz et al., 2018) in biological duplicates. 100,000 ESCs were crosslinked in 1% formaldehyde and incubated for 10 min at room temperature with rotation (20 rpm). The reaction was quenched by adding glycine (0.2 M) and incubating for 5 min at room temperature with gentle rotation (20 rpm). Cells were washed three times with 1 ml of cold PBS (centrifuged at 300 g for 5 min at 4°C) and then gently resuspended in 250 μl of ice-cold *in situ* Hi-C buffer (10 mM Tris-Cl pH 8.0, 10 mM NaCl, 0.2% IGEPAL CA-630, cOmplete Ultra protease inhibitors) and incubated on ice for 15 min. Samples were then centrifuged and resuspended in 250 µl of *in situ* Hi-C buffer. Cells were centrifuged (13,000 g for 5 min at 4°C) and resuspended in 250 µl ice-cold 10x NEB2 buffer. Nuclei were centrifuged (13,000 g for 5 min at 4°C) and permeabilised by resuspending them in 50 µl of 0.4% SDS and incubating at 65°C for 10 min. SDS was quenched by adding 25 µl of 10% Triton X-100 and 145 µl of nuclease-free water and incubated at 37°C for 45 min with shaking (650 rpm). Chromatin was digested by adding 100 U of MboI in 20 µl of 10x NEB2.1 buffer for 90 min at 37°C with rotation. MboI was heat-inactivated at 62°C for 20 min. The overhangs generated by the restriction enzyme were filled-in by adding a mix of 0.4 mM biotin-14-dATP (Life Technologies), 10 mM dCTP/dGTP/dTTP (0.75 µl of each dinucleotide), and 5 U/µl DNA polymerase I Klenow (8 µl; New England Biolabs), and incubated for 90 min at 37°C with rotation. DNA fragments were ligated in nuclease-free water (657 µl), 10x T4 DNA ligase buffer (120 µl), 10% Triton X-100 (100 µl), 20 mg/mL BSA (12 µl) and 5 Weiss U/µl T4 DNA ligase (5 µl in two instalments; Thermo Fisher) by incubating 4 h at 20°C with gentle rotation. Nuclei were centrifuged (2,500 g for 5 min at room temperature) and resuspended in 500 µl extraction buffer. Protein was digested with 20 µl of 20 mg/mL Proteinase K (Applichem), for 30 min at 55°C with shaking (1,000 rpm). 130 μL of 5M NaCl was added followed by overnight incubation at 65°C with shaking (1,000 rpm). Phenol-Chloroform-Isoamyl alcohol (25:24:1; Sigma-aldrich) extracted DNA was resuspended in 30 µl of 10mM Tris pH 8.0 (Applichem) and incubated for 15 min at 37°C with 10 mg/ml RNase A (1 µl; Applichem). In order to remove biotin from unligated fragments, DNA samples were incubated at 20°C for 4h without rotation in a mix of 10 µl of 10x NEB2 buffer (New England Biolabs), 1 mM of a dNTPs mix (10 µl), 20 mg/mL BSA (0.5 µl), 3 U/µl T4 DNA polymerase (5 µl; New England Biolabs) and nuclease-free water (up to 100 µl). Samples were sheared using a Covaris S220 instrument (2 cycles, each 50 sec, 10% duty, 4 intensity, 200 cycles/burst). Biotinylated fragments were pulled down using Dynabeads MyOne Streptavidin C1 beads. Libraries were end repaired on beads using the NEBNext Ultra End Repair module (New England Biolabs) and washed twice on 1x B&W (10 mM Tris-Cl pH 7.4, 1 mM EDTA, 2 M NaCl) + 0.1% Triton X-100, resuspended in 50 µl and transferred to a 1.5 mL tube. Adaptors for Illumina sequencing was added using the NEBNext® Ultra™ dA-Tailing module (New England Biolabs). Final amplification of the libraries was done in 4 parallel reactions per sample as follows: 10 µl of the bead-bound libraries, 25 µl of 2x NEBNext Ultra II Q5 Master Mix, 5 µl of 10 µM Universal PCR primer, 5 µl of 10 µM Indexed PCR primer and 10 µl of nuclease-free water.

Samples were individually barcoded and amplified for 10 (Tir1+Aux_Batch1, Ring1B+Aux_Batch1, Scc1+Aux_Batch1), 12 (Ring1B+Aux_Batch3) or 14 (Tir1+Aux_Batch3, Scc1+Aux_Batch3) cycles following the program: 98°C for 1 min, (98°C for 10 s, 65°C for 75 s, ramping 1.50°C/s) repeated 10-14 times, 65°C for 5 min, 4°C hold.

The four reactions were combined into one tube and size-selected using Ampure XP beads (Beckman Coulter). Final Hi-C libraries were quantified using Qubit dsDNA HS assay kit and a DNA HS kit on a 2100 Bioanalyzer (Agilent). Libraries were first pooled and shallow sequenced on an Illumina MiSeq (2×84bp paired-end; MiSeq reagent kit v3-150 cycles) to assess library quality. They were then sequenced on an Illumina NextSeq (2×80 bp paired-end; NextSeq 500/550 High Output kit v2-150 cycles).

### Hi-C Analysis

For each library, paired-end reads were independently mapped against the mm10 reference genome (UCSC) using Bowtie2 in ‘--very-sensitive’ mode. Unmapped reads were truncated by 8bp and realigned iteratively, until a valid alignment could be found or the truncated read was shorter than 30bp. Only uniquely mapping reads with a mapping quality (MAPQ)>=30 were kept in the downstream analysis. Biopython “Restriction” module was then used to compute predicted restriction fragments. Uniquely mapped reads were assigned to fragments, fragments to pairs and pairs filtered for self-ligated fragments, PCR duplicates, read pairs mapping further than 5 kb from the nearest restriction site, and for uninformative ligation products (Cournac et al., 2012). The genome was binned at 10 kb resolution, and Hi-C matrices were built by counting the number of valid fragment pairs per bin. Bins with less than 10% of the median number of fragments per bin were masked before the matrix was normalised using KR matrix balancing per chromosome (Knight and Ruiz, 2012).

### Observed/expected (OE) Hi-C matrix generation

Expected Hi-C contact values were obtained by calculating the average contact intensity for all loci with the same distance. The normalized Hi-C matrix is then transformed into an observed/expected (O/E) matrix by dividing each normalized observed by its corresponding expected value at that distance. O/E matrix generation was performed for each chromosome separately.

### A/B compartment quantification

A/B compartment calculation was done following a previously described procedure (Lieberman-Aiden:2009; Flyamer et al., 2017). Briefly, O/E matrices for each chromosome at 500 kb resolution were transformed into a correlation matrix by calculating the Pearson correlation of row i and column j for each (i, j). The first eigenvector of the correlation matrix forms the compartment vector. To ensure that positive values indicate the A (active) compartment and negative values the B (inactive) compartment, we used GC content as a proxy: if the average GC content of regions with negative entries is higher than that of regions with positive entries, the eigenvector sign is inverted. Absolute intra-chromosomal correlation values were compared between conditions as a measurement of compartmentalisation.

### Hi-C peak calling

SCC1-AID peaks were called in 100kb resolution matrices using an in-house, CPU implementation of HiCCUPs (Rao et al., 2014). Enrichment and FDR values for each pixel were obtained as described (Rao et al., 2014). Peaks must (i) have a minimum of 2.25-fold enrichment over the donut neighbourhood, (ii) have an FDR≤0.05 in the donut neighborhood, (iii) have an FDR≤0.1 in the remaining the neighbourhoods, and (iv) have a minimum observed value of 29 contacts in the peak centre. The robustness of these specific values has been confirmed visually for a large number of regions in order to minimise false-positives.

### Aggregate Hi-C feature analysis (TADs, peaks and A/B compartments)

Published ESC TAD intervals were used for aggregate TAD analysis (Bonev et al., 2017). For calculating the aggregate TAD and peaks, subsets of the O/E matrices were extracted and averaged to obtain the output sub-matrices. Sub-matrices of different sizes were interpolated using “imresize” with the “nearest” setting from the Scipy Python package. Using the TAD and peak calls for each of the groups (see the above section “Average Hi-C feature analysis” for parameter details). The aggregate analysis of the O/E matrices were calculated at 10kb resolution for TADs (Flyamer et al., 2017) and at 10 kb resolution for peaks.

### ChIP-Seq read enrichment quantitation at Hi-C peaks

Datasets in Supplementary Table S1 were processed using the standard pipeline in the lab (see cChIP-Seq read processing below). Pileups were built using MACS2 and the obtained bedgraph files were used to quantify read count enrichments. Read count enrichments were quantified separately for source and sink of each interaction using the function annotatePeaks.pl from HOMER (Heinz et al., 2010) with the options –size given-raw. For each peak, an average enrichment was quantified using the mean between source and sink. This was repeated for 1000 distance- and chromosome-matched random source-sink pairs. Fold enrichment was quantified by dividing observed enrichment by the mean enrichment at random source-sink pairs.

### Capture-C

#### Capture-C library generation

Capture-C libraries were prepared as described previously (Davies et al., 2016). 10^7^ mouse ES cells were trypsinized, collected in 50ml falcon tubes in 9.3ml media and crosslinked with 1.25 ml 16% formaldehyde for 10 min at room temperature. Cells were quenched with 1.5ml 1 M glycine, washed with PBS and lysed for 20 min at 4°C while rotating (lysis buffer: 10 mM Tris pH 8, 10 mM NaCl, 0.2% NP-40, supplemented with complete proteinase inhibitors) prior to snap freezing in 1 ml lysis buffer at −80°C. Lysates were then thawed on ice, pelleted and resuspended in 650 µl 1×DpnII buffer (NEB). Three 1.5ml tubes with 200 µl lysate each were treated in parallel with SDS (0.28% final concentration, 1 h, 37°C, interval shaking 500rpm, 30s on/30s off), quenched with trypsin (1.67%, 1h at 37°C, interval shaking 500rpm, 30s on/30sec off) and subjected to a 24 h digestion with 3×10 µl recombinant DpnII (37°C, interval shaking 500rpm, 30s on/30s off). Each chromatin aliquot was independently ligated with 8 µl T4 Ligase (240 U) in a volume of 1440 µl (20 h at 16°C). Following this, the nuclei containing ligated chromatin were pelleted, reverse-crosslinked and the ligated DNA was phenol-chloroform purified. The sample was resuspended in 300 µl water and sonicated 13x (Bioruptor Pico, 30s on, 30s off) or until a fragment size of approximately 200 bp was reached. Fragments were size selected using AmpureX beads (Beckman Coulter, selection ratios: 0.85x / 0.4x) and the correct size was assessed by Bioanalyzer. 2× 1-5 µg of DNA were adaptor ligated and indexed using the NEBNext DNA library Prep Reagent Set (New England Biolabs: E6040S/L) and NEBNext Multiplex Oligos for Illumina Primer sets 1 (New England) and 2 (New England). The libraries were amplified 7× using Herculase II Fusion Polymerase kit (Agilent).

#### Capture-C hybridization and sequencing

5’ biotinylated probes were designed using the online tool by the Hughes lab (CapSequm) to be 70-120bp long and two probes for each promoter of interest. The probes were pooled at 2.9nM each. Samples were captured twice and hybridizations were carried out for 72h and for 24h for the first and the second captures, respectively. To even out capture differences between tubes, libraries were pooled prior to hybridization. For Control, SCC1^DEG^, RING1B^DEG^ and SCC1^DEG^ RING1B^DEG^, 1.5µg of each replicate was individually hybridized and then pooled for the second round of hybridization. CTCF +/-AUX were multiplexed prior to the first capture at 2 µg each. Hybridization was carried out using Nimblegen SeqCap (Roche, Nimblegen SeqCap EZ HE-oligo kit A, Nimblegen SeqCap EZ HE-oligo kit B, Nimblegen SeqCap EZ Accessory kit v2, Nimblegen SeqCap EZ Hybridisation and wash kit) following manufacturer’s instructions for 72 h followed by a 24 h hybridization (double Capture). The captured library molarity was quantified by qPCR using SensiMix SYBR (Bioline, UK) and KAPA Illumina DNA standards (Roche) and sequenced on Illumina NextSeq 500 platform for three biological replicates.

#### Capture-C data analysis

Fastq files were aligned to mm10 genome and filtered using HiCUP (v0.5.7) (Wingett et al., 2015) and Bowtie 2 (Langmead et al., 2009) with the settings of 100bp-800bp for fragment sizes. Paired bam files were then processed using the Bioconductor package Chicago (Cairns et al., 2016) (Version: 1.0.4) according to the Chicago Vignette using the inbuilt mESC-2reps weight settings. Interaction “peaks” were called based on Chicago scores >=5 and interaction peaks closer than 10 fragments in distance were combined to one peak. Weighted average read counts were extracted from the ChicagoData objects. For visualization in line plots, for each DpnII fragment, percentage reads per promoter (PRPP) was calculated for each sample to normalize the read counts. Briefly, read counts were divided by the total coverage of reads aligned to captured promoters in the sample, multiplied by the amount of promoters captured and then multiplied by 100 to obtain % reads per promoter captured for each DpnII restriction fragment (PRPP = N / cov * nprom * 100). For display purposes reads were then multiplied by 1000 for Figures 3-5 (mPRPP). For aggregate peak analysis (Fig. 3-4) significantly enriched interactions were determined using Chicago default threshold of score >= 5 at the level of individual DpnII fragments. Because peaks between polycomb occupied sites are larger than the average DpnII fragment, interactions with <10 DpnII fragments distance were merged to one peak. Peak summits were then defined as the local maximum in the Control sample (if peaks were present in this sample) or in the sample in which they were present. In order to make interactions at different distances comparable, all samples were then normalized to PRPPs at the peak summit in Control. For aggregate analyses in Figures 3- 4 only interactions between polycomb target gene promoters and a stringent set of RING1B peaks (Fursova et al., 2019, in press) were considered.

### Calibrated RNA-Seq and ChIP-Seq

#### Calibrated total RNA-seq (cRNA-seq)

To prepare RNA for cRNA-seq, 5 million mouse ESCs (SCC1-AID +/-Auxin) were mixed with 2 million Drosophila SG4 cells. Total RNA was extracted using RNeasy Mini Kit (QIAGEN) according to the manufacturer’s protocol, followed by treatment with the TURBO DNA-free Kit (ThermoScientific). Quality of RNA was assessed using 2100 Bioanalyzer RNA 6000 Pico kit (Agilent). To construct libraries, for each sample RNA was first depleted of rRNA using the NEBNext rRNA Depletion kit (NEB). RNA-seq libraries were then prepared from 200 ng of RNA using the NEBNext Ultra II Directional RNA-seq kit (NEB). To quantitate the consistency of spike-in cell mixing for each individual sample, genomic DNA was isolated from a small aliquot of mixed mouse and fly cells using Quick-DNA Miniprep kit (Zymo Research) according to the manufacturer’s protocol. Libraries from 50 ng of genomic DNA were constructed using NEBNext Ultra II FS DNA Library Prep Kit (NEB), following manufacturer’s guidelines. NEBNext Multiplex Oligos were used for indexing libraries. The average size of all libraries was analysed using the 2100 Bioanalyzer High Sensitivity DNA Kit (Agilent) and the libraries concentration was measured by qPCR using SensiMix SYBR (Bioline, UK) and KAPA Illumina DNA standards (Roche). cRNA-seq and gDNA-seq libraries were sequenced as 80 bp paired-end reads on the Illumina NextSeq 500 platform for four independent biological replicates.

#### Calibrated ChIP-Seq

50 million Control or SCC1^DEG^ mESCs were mixed with 500,000 HEK293 cells before fixation. Cells were fixed for 10 minutes in 1% formaldehyde at room temperature. Formaldehyde was quenched by the addition of glycine to a final concentration of 125 µM. All subsequent steps were as previously described (King and Klose, 2017). Libraries were sequenced for three biological replicates.

#### Massively parallel sequencing, data processing and normalisation

For cRNA-seq, to filter out reads mapping to rDNA fragments, paired-end reads were aligned using Bowtie 2 (with “--very-fast”, “--no-mixed” and “--no-discordant” options) against the concatenated mm10 and dm6 rRNA genomic sequence (GenBank: BK000964.3 and M21017.1). All unmapped reads from this step were then aligned against the genome sequence of concatenated mm10 and dm6 genomes using the STAR aligner (Dobin et al., 2013). Finally, reads that failed to map using STAR were additionally aligned against the mm10+dm6 concatenated genome using Bowtie 2 (with “--sensitive-local”, “--no-mixed” and “--no-discordant” options). Uniquely aligned reads from the last two steps were combined for further analysis. PCR duplicates were removed using SAMTools. For cChIP-Seq, we aligned paired-end reads to a concatenated mouse and human genome (mm10+hg19) using Bowtie2 with “--no-mixed” and “--no-discordant” options and SAMBAMBA (Tarasov et al., 2015) was used to filter out PCR duplicates. The mean and standard deviation of the insert size was calculated using Picard tools. To visualise gene expression changes, uniquely aligned mouse reads were normalised using drosophila (or human for cChIP-Seq) spike-in as described previously (Hu et al., 2015). Briefly, mm10 reads were randomly subsampled based on the total number of dm6 (or hg19) reads in each sample. To account for any minor variations in spike-in cell mixing between replicates, the subsampling factors were additionally corrected using the ratio of dm6 (or hg19)/mm10 total read counts in corresponding gDNA-seq samples. Genome coverage tracks were then generated with genomeCoverageBed from BEDTools (Quinlan, 2014) and visualised using the UCSC genome browser (Kent et al., 2002).

#### Read count quantitation and differential gene expression analysis

For differential gene expression analysis, a custom-built non-redundant mm10 gene set was used to obtain read counts from original bam files prior to spike-in normalisation using a custom Perl script. To generate the non-redundant mm10 gene set (n = 20,633), mm10 refGene genes were filtered to remove very short genes with poor sequence mappability and highly similar transcripts. To identify significant changes in gene expression following auxin treatment, a custom R script utilising DESeq2 package was used (Love et al., 2014). To incorporate spike-in calibration, raw mm10 read counts were normalised using DESeq2 size factors which were calculated based on the read counts for the set of unique dm6 refGene genes as previously described (Taruttis et al., 2017). Prior to quantitation, Drosophila reads were pre-normalised using the actual spike-in ratio (dm6/mm10) which was derived from a corresponding gDNA-seq sample. A threshold of p-adj < 0.05 and fold change > 1.5 was used to determine significant changes in gene expression. For visualisation normalised read counts were extracted from the DESeq2 table and used to quantify RPKM. These were log2 transformed after addition of a pseudocount of 0.01. Replicate correlations were calculated using the R Bioconductor function cor(method=’spearman’) from the package stats and were >0.99 throughout. Given the high reproducibility, DEseq2 normalized read counts for the replicates were pooled, RPKM normalized and log2 transformed as described above for visualization in Figure 6.

#### Read count quantitation and enrichment analysis for cChIP-Seq

For cChIP-Seq analysis, reads were quantified in a custom set of RING1B peaks. Paired reads were quantified using the function summarizeOverlaps() from the R Bioconductor package “GenomicFeatures” (Lawrence et al., 2013) with the option mode=”Union”. A pseudocount of 8 was added prior to log10 transformation. Replicates were compared using the cor(method=’spearman’) function from the R Bioconductor stats package and were >0.99. For pooled read counts, BAM files were merged using samtools and reads were quantified from merged BAM files using the procedure described above. Metaprofiles were obtained using the computeMatrix and plotHeatmap functions from deepTools suite (Ramírez et al., 2015).

#### RASER (Resolution After Single-strand Exonuclease Resection)-FISH

RASER-FISH was conducted as previously described (Brown et al., 2018) with minor changes. Briefly, cells were labelled for 24 h with BrdU/BrdC mix (3:1) at final conc. of 10 µM, with auxin added at 500 µM for the final 6 h. Cells were fixed in 4% PFA (vol/vol) for 15 min and permeabilised in 0.2% Triton X-100 (vol/vol) for 10 min. Cells were then stained with DAPI (0.5 µg/mL in PBS), exposed to 254 nm wavelength UV light for 15 min, then treated with Exonuclease III (NEB) at 5 U/μL at 37°C for 15 min. Labelled probes (100 ng each) were denatured in hybridization mix at 90°C for 5 min and pre-annealed at 37°C for 10 min. Coverslips were hybridized with prepared probes at 37°C overnight. Following hybridization, coverslips were washed for 30 min twice in 2× SSC at 37°C, once in 1xSSC at RT. Coverslips were blocked in 3% BSA (wt/vol) and digoxigenin was detected with sheep anti-digoxigenin FITC 1/50 (Roche, 11207741910) followed by rabbit anti–sheep FITC 1/100 (Vector Laboratories, FI-6000). Coverslips were stained with DAPI (0.5 μg/mL in PBS), washed with PBS and mounted Vectashield (Vector Laboratories).

#### Probes and nick-translation labelling

Fosmid probes WIBR1-0935O10 (Nkx2.3 mm9; chr19; 43,659,682-43,698,592), WIBR1-1122P14 (Pax2 mm9; chr19; 44,809,035-44,851,675), WIBR1-1125H10 (Dlx2 mm9; chr2: 71374041-71411685), WIBR1-2777G14 (HoxD10 mm9; chr2: 74511607-74550498) were obtained from BACPAC Resources Center (Children’s Hospital Oakland Research Institute; [https://bacpacresources.org/]). Probes were labelled for use in FISH by nick translation as follows: prior to nick translation, 1 μg DNA was treated with RNase (0.02 U) (Sigma), for 30 min at 37°C, nick translation was carried out at 16 °C for 1 h in the following reaction mixture; 50 mM Tris-HCl, 5 mM MgCl_2_, 2.5 μg BSA, 10 mM β-mercaptoethanol, 50 mM dAGC, 20 μM hapten/fluor [digoxigenin-11-dUTP (Sigma); Cy3 dUTP (GE Healthcare)], 15 U recombinant DNase1 (Sigma) and 10 U DNA polymerase I (NEB), made up to a final volume of 50 μl with H_2_0.

#### Imaging Equipment and Settings

Widefield fluorescence imaging was performed at 20°C on a DeltaVision Elite system (Applied Precision) equipped with a 100 × /1.40 NA UPLSAPO oil immersion objective (Olympus), a CoolSnap HQ2 CCD camera (Photometrics), DAPI (excitation 390/18; emission 435/40), FITC (excitation 475/28; emission 525/45) and TRITC (excitation 542/27; emission 593/45) filters. 12-bit image stacks were acquired with a z-step of 150 nm giving a voxel size of 64.5 nm × 64.5 nm × 150 nm. Image restoration was carried out using Huygens deconvolution Classic Maximum Likelihood Estimation (Scientific Volume Imaging B.V.).

#### Image Analysis

As previously described (Brown et al., 2018). Briefly, 3D distance measurements were made using an in-house script in ImageJ [https://imagej.net/]. As a pre-processing step image regions were chromatically corrected to align the green and the red channel images. Parameters for the chromatic correction were calculated through taking measurements from images of 0.1 μm TetraSpeck® (Molecular Probes®) and calculating the apparent offset between images in each colour channel. Cells were only selected for analysis where there was no hint of replicated signal. Signal pairs were manually identified whereupon a 20 × 20 pixel and 7-15 z-step sub-volume was automatically generated centered on the identified location. In each identified region, thresholding was applied to segment the foci. Firstly, the image region was saturated beyond the top 96.5 % intensity level, to reduce the effect of noisy pixels, and then the threshold was calculated as being 90 % of the maximum intensity value of the processed image. This was repeated for both green and red channels. Once segmented, signal centroid positions were mathematically calculated and the inter-centroid 3D distance measurement was output along with a .png image for visual inspection.

#### Contact probability threshold calculation

To assess what proportion of our inter-probe distance measurements might be considered as co-incident we applied the following rationale, which is as described (Cattoni et al., 2017). To measure the error in colocalisation precision within a realistic, non-ideal experimental situation, we labelled and hybridised the same fosmid probe with both digoxygenin (detected with FITC) and Cy3 in cells, as per the experimental conditions. The distance range measured between those two colours shows the colocalisation precision error of 73nm ± 38 nm (mean ± SD) in our experimental system. From this we conservatively assume that two probes have 99% chance of co-localisation if their separating distances are less than 187 nm (i.e. mean + 3xSD).

#### Antibodies

RAD21, 1:1000 (Abcam, ab154769), RING1B (western) 1:1000 (Klose Lab), RING1B (ChIP-Seq) 1:1000 (Cell Signalling, 5694), CTCF 1:1000 (Abcam, ab70303) and TUBULIN 1:500 (Abcam, ab6046).

---

**Supplementary Table 1–.**
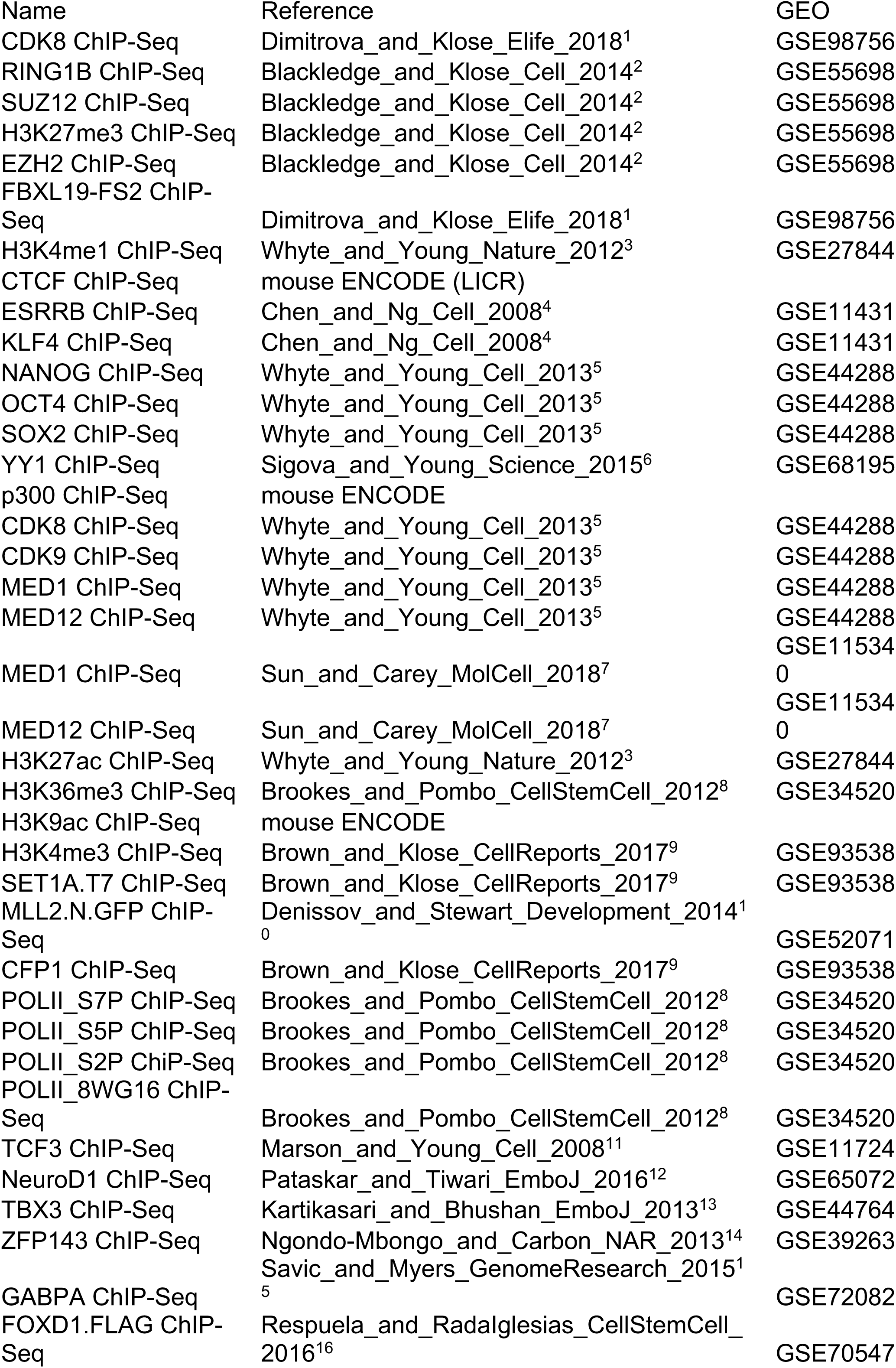
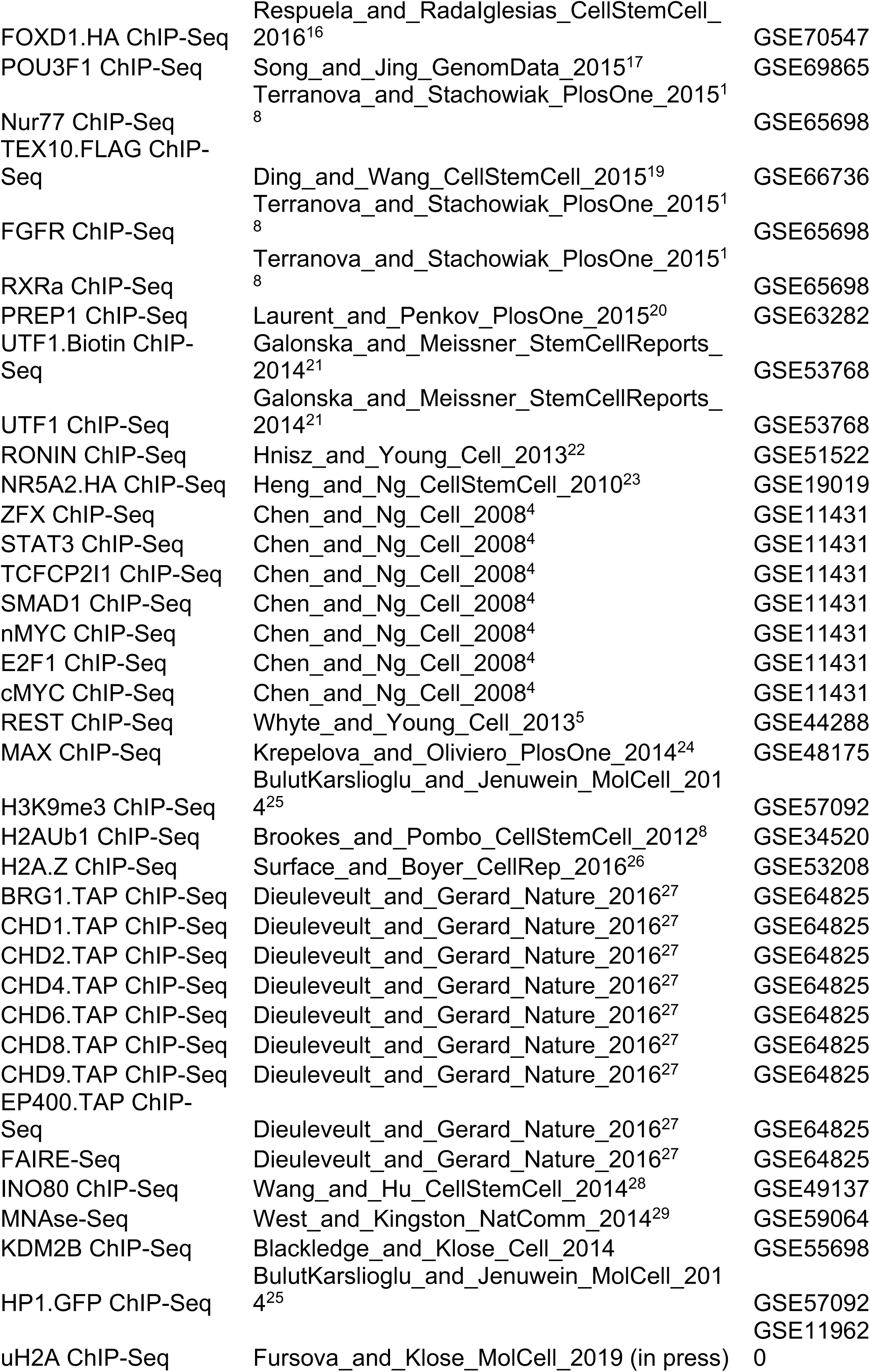
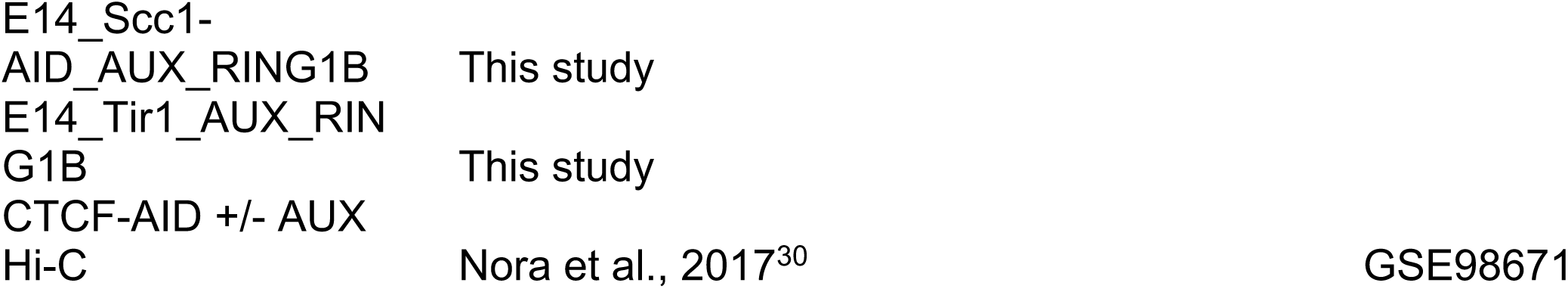
Dataset analysed in this study

